# Auxin-mediated stress relaxation in pericycle and endoderm remodelling drive lateral root initiation

**DOI:** 10.1101/2023.09.18.558377

**Authors:** João R. D. Ramos, Blanca Jazmin Reyes-Hernández, Karen Alim, Alexis Maizel

## Abstract

Plant development relies on the precise coordination of cell growth, which is influenced by the mechanical constraints imposed by rigid cell walls. The hormone auxin plays a crucial role in regulating this growth by altering the mechanical properties of cell walls. During the post-embryonic formation of lateral roots, pericycle cells deep within the main root are triggered by auxin to resume growth and divide to form a new root. This growth involves a complex interplay between auxin, growth, and the resolution of mechanical conflicts with the overlying endodermis. However, the exact mechanisms by which this coordination is achieved are still unknown. Here, we propose a model that integrates tissue mechanics and auxin transport, revealing a connection between the auxin-induced relaxation of mechanical stress in the pericycle and auxin signalling in the endodermis. We show that the endodermis initially limits the growth of pericycle cells, resulting in a modest initial expansion. However, the associated stress relaxation is sufficient to redirect auxin to the overlying endodermis, which then actively accommodates the growth, allowing for the subsequent development of the lateral root. Our model uncovers that increased pericycle turgor and decreased endodermal resistance licence expansion of the pericycle and how the topology of the endodermis influences the formation of the new root. These findings highlight the interconnected relationship between mechanics and auxin flow during lateral root initiation, emphasizing the vital role of the endodermis in shaping root development through mechanotransduction and auxin signalling.

## Introduction

The rigidity of plant cell walls imposes precise coordination of growth among cells, where spatial variations in growth generate mechanical stresses serving as instructive cues. This feedback mechanism links tissue geometry to gene expression (1, 2). Auxin, a pivotal plant hormone involved in cell growth and differentiation (3–9), plays a central role in growth regulation by modifying cell wall properties (10, 11). Its distribution relies on polar transport regulated by influx and efflux carriers (12–14). Efflux carrier polarization, particularly the PIN protein family, directs auxin flow, regulating plant growth and development (15–17). In the shoot meristem, PIN proteins have also been found to possess mechanical-sensing properties (18, 19), contributing to the complex interplay between auxin-mediated cell wall softening, tissue mechanics, and PIN-mediated auxin transport, ultimately shaping developmental auxin patterns (18, 20). Specifically, efflux carrier distribution correlates with the microtubule orientation and both are affected by stress orientation.

A prime example of coordinated cell and tissue growth is the formation of lateral roots (LRs) (21–23). In *Arabidopsis thaliana*, LRs originate from pericycle cells beneath the endodermis (24, 25). These Xylem pole pericycle (XPP) cells, typically in pairs, become activated through auxin accumulation, driven by local synthesis (26), lateral auxin diversion (27–29), and endodermal-to-pericycle auxin reflux (30) (Fig. 1 A). Activated XPP cells expand radially, pressing against the overlying endodermis (31). Nuclei migrate, triggering asymmetric cell division and lateral root primordium formation (21, 23, 32). Notably, this radial expansion is more pronounced at shared cell walls (31, 33), necessitating cortical microtubule (CMT) rearrangements in founder cells (33). Reciprocal auxin flows between the endodermis and the pericycle, accompanying lateral root initiation. Just before the founder cell radial expansion, active auxin transport via PIN3 from the endodermis to the pericycle aids accumulation in the founder cells (30). Upon radial expansion, the change in endodermis shape depends on auxin signalling in the endodermis, which implies auxin accumulation in the endodermis (31, 34). It is unknown whether this auxin originates from the pericycle and is part of the signal instructing the endodermis of the growth of the pericycle. It is also unknown which modification(s) of the mechanical properties of the pericycle and/or endodermis, such as turgor difference, licences the radial expansion.

**Fig. 1.**
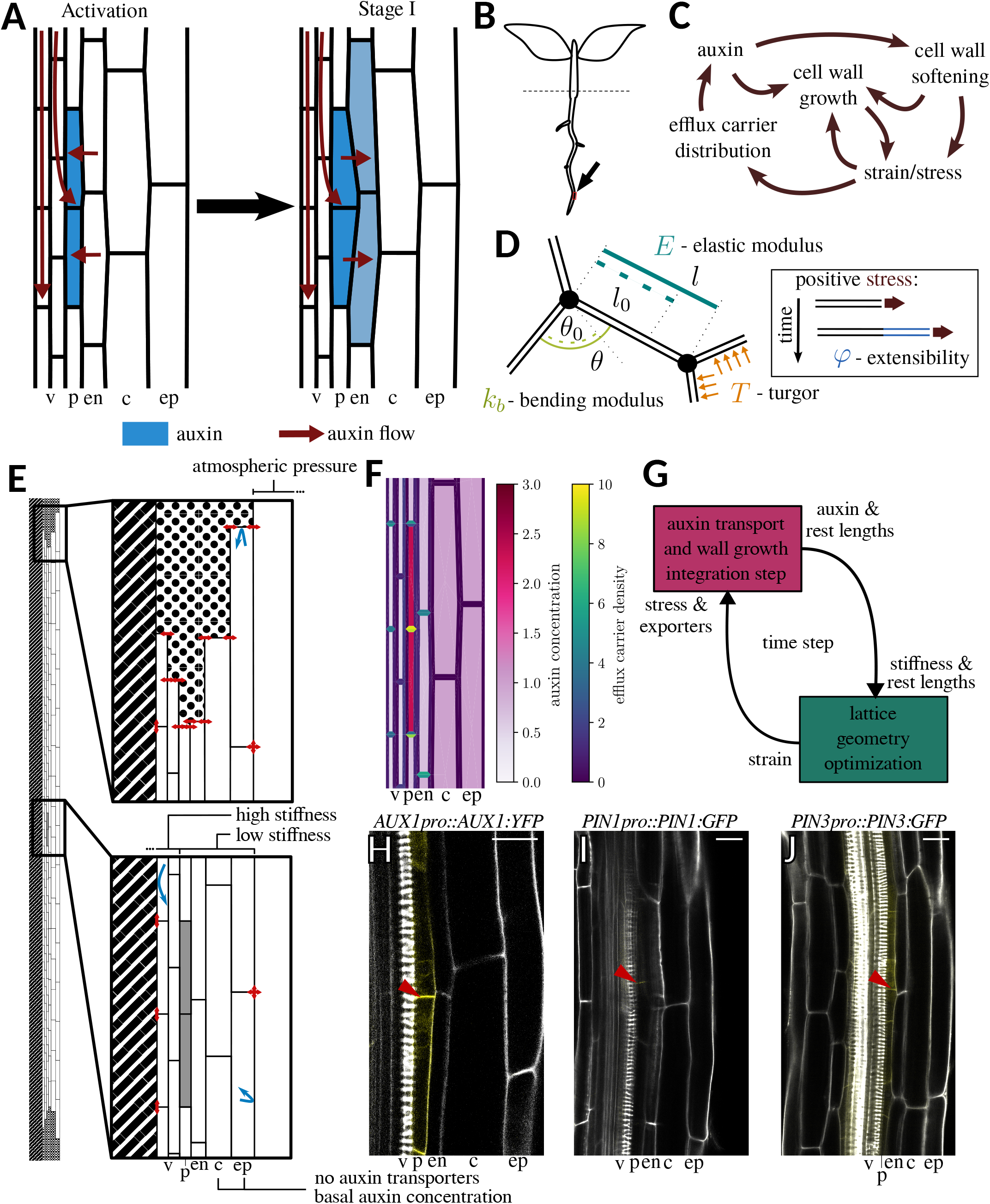
Biological context and modelling framework. (A) Activation of pericycle (p) before lateral root initiation results from the accumulation of auxin in these cells by local auxin synthesis, redirection of the rootward auxin flow in the vasculature (v) and from the endodermis (en). Upon activation, the pericycle cells with high auxin acquire a trapezoidal shape, and the endodermis accommodates this morphological change by shrinking. This accommodation of the endodermis requires the endodermis cells to perceive auxin. The diagram visualises the cellular template used in our mechanical vertex model, including the cortex (c) and epidermis (ep). (B) Diagram of an Arabidopsis seedling showing the location (arrow) we model here. (C) The interplay between auxin accumulation and cell wall mechanics: Auxin induces growth by altering the properties of the cell wall, which modifies the patterns of mechanical stress. Efflux carriers of auxin respond to mechanical stress perturbations, creating a mechanical feedback loop that affects the auxin concentration and flow pattern. (D) In our model, cell-cell interfaces in their entirety, including primary and secondary cell walls, membrane and middle lamella, are modelled as viscous rods that resist longitudinal deformations with an elastic modulus, *E*, and bending at the junctions with bending modulus *k*_*b*_. Turgor, *T*, generates stresses to which the walls respond by growing according to the extensibility, *φ*, of the respective viscous rod. (E) Lattice used for simulations. The left-most vertices form a boundary representing the inside of the root, and the right-most vertices define the plant’s boundary, which faces a container of lower pressure. The innermost vertices can only move parallel to this boundary, whereas the vertices on the outer boundary are free to move in general (lower zoom in, red arrows). On the upper and lower boundaries, we emulate the effects of having neighbours pushing back against the tissue with the same constraints, leading only to horizontal displacements and resulting in vertical confinement (upper zoom in, red arrows). From the pericycle layer, two cells are selected as primed (lower zoom-in, grey). Auxin is reflected at the boundaries (blue arrows). (F) After a few iterations, depletion around the primed cells is visible, and auxin progressively accumulates in the two primed XPP cells. Most efflux carriers bind from one cell to the other. (G) Schematic of the order of operations during a time step of our model along with the inputs and outputs of each major section. (H, I, J) Experimental results of bound carriers during stage 0 LR initiation (arrowhead denotes the shared wall between activated XPP cells). (H) The influx carrier AUX1 is predominantly expressed in these two cells. (I, J) Efflux carriers are only detected in and around the primed XPP cells and the vasculature. Scale bar: 20 *µ*m.

Here, building on the conceptual framework established in the shoot apical meristem, we model the interplay between growth mechanics and auxin flow during LR initiation (Fig. 1 B). Our model combines a vertex model for tissue mechanics and a compartment model for auxin transport (20). The growth of the shared anticlinal wall between adjacent XPP cells is constrained, allowing only modest radial swelling. However, auxin-induced stress relaxation in this wall relocates auxin efflux carriers to the periclinal walls, redirecting auxin flow to the endodermis and triggering an auxindependent response. Empirical observations support this prediction. Furthermore, we demonstrate that endodermal wall properties and arrangement limit the radial expansion of activated XPP cells. These findings propose a biphasic radial growth model for activated XPP cells, initially constrained by the endodermis, enabling auxin redirection to the endodermis. This, in turn, triggers an auxin-dependent response, facilitating radial expansion of activated XPP cells.

## Materials and Methods

We previously developed a two-dimensional vertex model that integrates a compartment model for auxin transport to explore the interaction between tissue mechanics and auxin patterns (20). This approach is similar to previous works on plant tissue mechanics using mass-spring models (35, 36), yet within a vertex model framework (37–39), and auxin transport models employed to study phyllotaxis (16, 18, 40, 41). Additionally, the viscous stress-based growth of cell walls is modelled similarly to (42, 43). Here, we describe the model used and its implementation. The symbols used are described upon their usage and in Table 1. For a more detailed and formal version, refer to the Supporting Material.

**Table 1.**
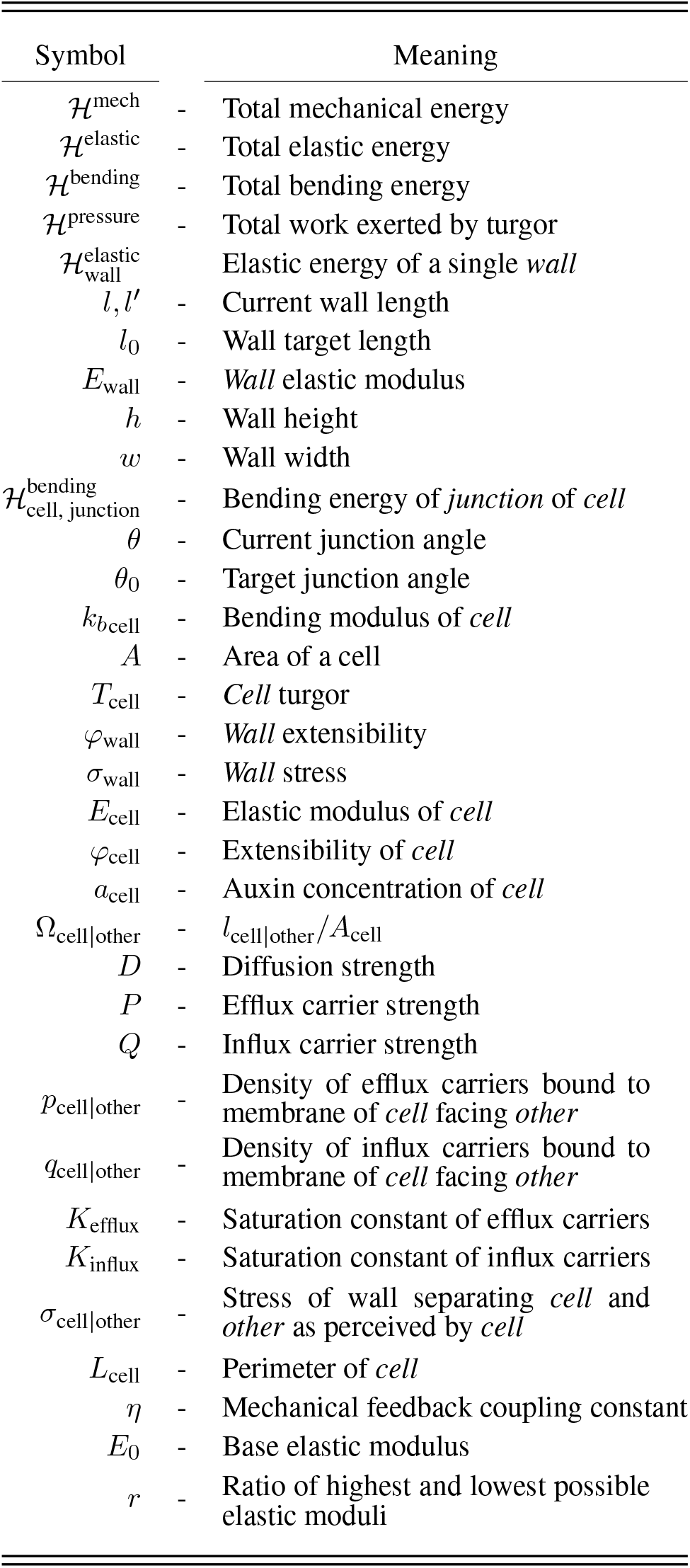
List of Symbols.

### Vertex model describes tissue mechanics

Cells are subject to hydrostatic pressure, and the interfaces between them exhibit resistance to compression, extension, and resistance to bending at junctions (Fig. 1 C). We minimize the total mechanical energy of the tissue, ℋ^mech^, decomposed as:

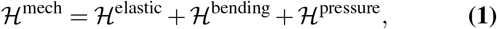

where ℋ^elastic^ represents the energy cost of longitudinal deformation (extension or compression) of the interface between cells, ℋ^bending^ the energy cost of changing the internal angle of the cell at one of its junctions, and ℋ^pressure^ the work done by cell turgor.

We describe the deformation of the interface by a simple elastic term of an elastic rigid one-dimensional rod as if it were a wall, similar to (35, 36). Here, the term wall refers to these rods as a proxy for the cell-cell interface, including primary and secondary cell walls, membrane and middle lamella. The total elastic energy, ℋ^elastic^, is the sum over all individual cell walls, each contributing the quantity

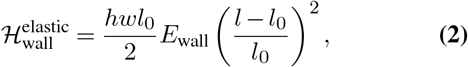

where *h* and *w* denote the height and width, *l*_0_ the rest length, *l* the current length, and *E*_wall_ the elastic modulus of the wall element in question. Note, the elastic modulus of a wall, *E*_wall_, will be a function of the auxin content of the surrounding cells.

We base our approach to wall bending energy on models often employed in the field of polymer physics (44–46) and assign a cost to change the angle, *θ*, concerning its rest value, *θ*_0_. Then, we account for each junction of each cell for the bending contribution to the mechanical energy,

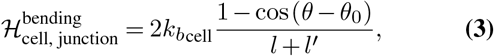

where *l* and *l*^*′*^ are the lengths of the walls whose intersection forms the junction in question, and 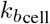 the bending modulus of the cell whose internal angle we are measuring. This allows us to readily assign a bending modulus following the cell’s internal processes and implicitly couple the contribution of several cells surrounding a particular junction.

The final term in Eq. 1 represents the effect of cell turgor, summed over all cells *T*_cell_. The work of internal hydrostatic pressure per cell of area, *A*, contributes

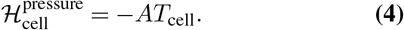

Therefore, given a set of rest lengths, rest angles, cell elastic moduli, bending moduli, and turgor pressures, we can compute the mechanical energy for a configuration of the tissue and by minimising the mechanical energy for the junction positions, we obtain the tissue configuration balancing all forces arising from the competing elastic, bending and pressure contributions.

### Growth as a viscous wall response

Cell walls grow when under positive strain/stress (47, 48). To model this behaviour, we describe the cell wall as a Maxwell material similar to (42, 47), by taking irreversible deformation into account. Because of how we discretize the system, we opt for the approach of Corson et al. (42). We incorporate cell wall growth into our model by defining the rate of change of the rest length, *l*_0_, of a particular wall with length *l*, elastic modulus *E*_wall_, and extensibility *φ*_wall_ (here defined as the reciprocal of viscosity) as

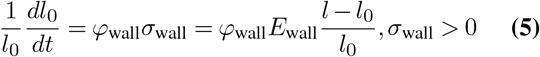

where *σ*_wall_ represents the longitudinal stress acting on the wall (Fig. S1 A). As mentioned before, the properties of each wall are a function of the processes taking place inside the surrounding cells. Therefore, we want to link the quantities *E*_wall_ and *φ*_wall_ to cell-specific counterparts, *E*_cell_ and *φ*_cell_. In a similar approach to (35), we associate elastic elements in parallel, resulting in *E*_wall_ being the average of *E*_cell_ of the two surrounding cells (Fig. S1 B). We also associate the viscous elements in parallel, assuming each viscous element is affected by the same stress as the corresponding elastic element (Fig. S1 B). If wall A|B separates cells A and B, then we obtain

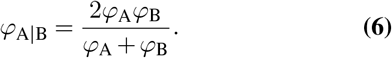

Interestingly, this coupling implies that extensibility, and, therefore, growth rate, is limited by the least extensible wall. This result can be obtained by inspecting the limit *φ*_*A*_ ≫ *φ*_*B*_. If the extensibilities are the same between the two cells, then so is the extensibility of the wall. Also, keeping a Maxwell material’s overall constitutive relation allows us to keep a single value of rest length, *l*_0_.

### Auxin transport as a compartment model

At the same time as we integrate wall growth, we also integrate auxin transport, which is described by a set of ordinary differential equations relating the quantity of auxin of a cell to the auxin quantity of the same cell and its neighbours in the previous time step. We assume auxin, being a small molecule, diffuses quickly within the cell and, therefore, we approximate auxin concentration as a constant value inside the cell. The auxin concentration within each cell is regulated by synthesis, influx, and efflux carriers, with efflux carriers responding to membrane stress. Higher cell membrane stress increases efflux carrier accumulation (19), facilitating auxin transport up a concentration gradient (18, 20, 35). Our approach here differs to from (20) in considering influx carriers and efflux carriers. Auxin concentration of a cell, *a*_cell_, evolves over time according to

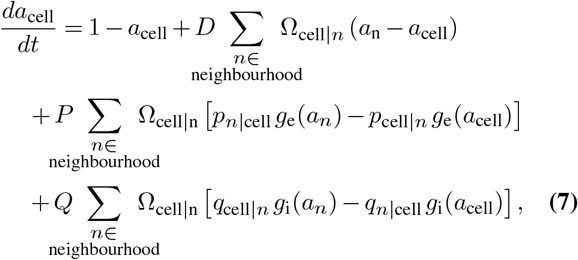

where *D, P*, and *Q* determine the strength of diffusion, active efflux, and active influx respectively, *p*_A|B_ means the density of efflux carriers bound to the membrane of cell A pumping auxin in the direction of cell B, *q*_|AB_ is the density of influx carriers bound to the membrane of cell A taking auxin from cell B, Ω _A| B_ = *l*_A|B_*/A*_A_, i.e., the length of wall separating cells A and B divided by the area of cell A, and, finally,

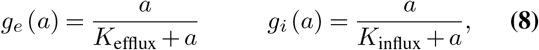

where *K*_efflux_ and *K*_influx_ define how quickly the activity of the respective carriers saturates with auxin concentration. The first and second terms in the right-hand-side of Eq. 7 stand for auxin synthesis and decay, the third for passive transport and the last two for active auxin efflux and influx, respectively. Note that the auxin transport equation has adimensional units of time and auxin concentration such that *a* = 1 is the base value cells would acquire by synthesis and decay alone and a decay rate equal to 1.

Although influx carrier density will be set according to experimental observations (Fig. 1 H), we model efflux carriers as being regulated by stress. The pattern formation mechanism proposed by (18) relies on having a differing value of membrane stress on each side of a single wall (18, 20), therefore we write down the stress perceived by cell A in contact with wall A B, of length *l* and rest length *l*_0_ as *σ*_A| B_>= *E*_A_ (*l*−*l*_0_) */l*_0_ ≠ *σ*_B|A_. Taking into account competition for the same pool of efflux carriers within a cell and modelling binding rate as a power law of stress, we obtain

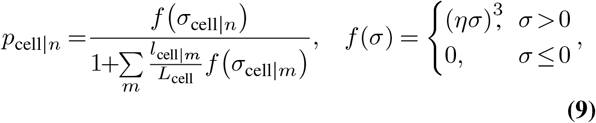

where *n* and *m* are cells in the corresponding neighbourhood, *L*_cell_ is the perimeter of the cell, and *η* is a coupling constant. Note that efflux carrier distribution is not dynamic but the steady-state result of a parsimonious carrier binding model (20).

### Auxin modulates cell wall mechanics

The effect of auxin on mechanical properties (10, 11) allows efflux carriers to sense auxin content and gives rise to a modulation of wall elastic modulus by auxin concentration written as(18, 20),

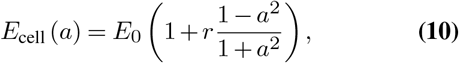

where *E*_0_ is the value of the elastic modulus at base auxin concentration, which we will call base elastic modulus for short, and *r* is the ratio between the maximum and minimum elastic modulus the tissue can exhibit in response to auxin. A change in auxin concentration leads to a change in efflux carrier distribution, resulting in up-the-gradient auxin flows. The bending modulus ought to be related to the elastic modulus in some way; however, due to the composition of the cell wall, it is difficult to tell what that relationship might look like, and thus, we vary them independently.

### Parametrization

Values for the different parameters were either obtained from the literature or imputed to ensure stable tissue behaviour. Details are provided in supplemental material.

### Implementation

Given an initial lattice of cells and the initial auxin concentration of each cell, as well as the initial rest length for each wall, along with the mechanical parameters *E*_cell_,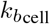, *T*_cell_, and target angles, *θ*_0_, we minimize mechanical energy, arriving at a new geometry. We do so while conserving auxin content before and after geometrical changes. With new wall lengths, *l*, we compute strain and stress. We then advance the state of rest lengths and auxin concentrations with Eq. 5 And Eq. 7. We, thus, arrive at a new set of values of auxin, and hence of *E*_cell_, and rest lengths for which our geometrical configuration is no longer at minimum energy. We iteratively repeat the above process until we reach the desired simulation time (Fig. 1 G). The simulation program was written in C++, using the Quad-Edge implementation by Paul Heckbert (49), and wrapped up in Python as a library with SWIG. The optimization routine used is a lowstorage-BFGS from the NLopt package (50). As the lattice deforms at each time step, we use a simple explicit forward Euler method for ODE integration. The NumPy (51) and Matplotlib (52) packages were used to read data and generate graphics.

### Plant material and growth

Seedlings of sC111 (*pUBQ10::GFP:PIP1*; *pGATA23::H2B:mcherry*; *pDR5v2::YFP:nls*) (33) and *pSHY2::nls:mVenus*; *pLBD16::mCherry:SYP122* (31) were used to monitor DR5v2 and SHY2/IAA3, respectively. Seeds were surface-sterilized with Ethanol 70% and SDS 0.05% for 5 min, rinsed once with Ethanol 99% and air-dried. After stratification at 4°C for 48 h, seedlings were grown vertically on square Petri dishes plates containing Murashige and Skoog 0.5× medium with a pH of 5.7, 0.8% agar and 1% sucrose, at 22°C with a photoperiod of 8/16 h light/dark condition. Four days after germination, lateral root primordium (LRP) induction through 5 h of gravistimulation (turned plates vertically, 180°) to detect early events of LRP formation was applied and to proceed with live imaging.

### Live Imaging and Microscopy

For imaging, the seedlings were transferred to a chambered cover glass as described in (53). Propidium Iodide (2 *µ*g*/*mL) was added when required. Confocal Laser-Scanning Microscopy (CLSM) was performed on a Leica SP8 confocal microscope with a 63×, NA = 1.2 water immersion objective. GFP, YFP and mVenus fluorescence were detected using the 488 nm and 514 nm excitation laser line and a detection range of 495-560 nm and 524-550 nm, respectively. mCherry and Propidium Iodide fluorescence was detected using the 568 nm excitation laser line and a detection range of 590-750 nm. Images were taken every 30 min after LRP induction for ∼12 h with a resolution of 1024 × 1024 pixels, 400 Hz speed and a line average of 4. Z stacks were performed using a z-step size of 0.5mm. Postprocessing of images (LUT, Bright, contrast and maximum intensity projections of 3-7 z-sections was performed in Fiji (http://fiji.sc/Fiji)).

## Results and disccussion

To simulate the interplay between tissue mechanics and auxin flow during lateral root initiation, we used a cellular lattice obtained from optical longitudinal sections of *Arabidopsis thaliana* roots, matching the *in vivo* aspect ratios. Our lattice has six cell layers representing vasculature, pericycle cells, endodermis, cortex, and epidermis (Fig.1 A, E). Although the model is run on an extended longitudinal section of the root (Fig. 1 B, E), we are mostly interested in a pair of activated XPP cells and their surroundings, and only this area is depicted.

Accumulation of auxin in root pericycle cells specifies them as lateral root founder cells (24). To ensure this auxin maximum is stable, we set auxin influx carrier density to be high, in agreement with experimental data (Fig. 1 H). In addition, we assign a higher elastic modulus to the vasculature to mimic the rigid behaviour of the xylem. This also aids auxin accumulation by biasing efflux carrier polarity from the vasculature to the pericycle. We initialise two neighbouring activated pericycle cells with double the amount of basal auxin concentration, reflecting the initiation process (32). To give these two pericycle cells the best chance to expand radially, we also set their bending modulus to *k*_*b*_ = 100 MPa.

The vertices facing the boundary towards the inside of the root are only allowed to move on the root-shoot axis. In contrast, vertices facing the outside of the root can move in any direction and tend to move outwards due to the lower pressure outside the plant, resulting in the sustained stress of anticlinal walls. We choose this pressure to be 0.1 MPa, which is approximately the value of atmospheric pressure. The simulated tissue is confined within a section of the root-shoot axis with a similar boundary condition to the vertices facing the boundary towards the inside of the root (Fig. 1 E). This means stress on periclinal walls is a free parameter we can control by setting appropriate initial wall rest lengths. Lastly, auxin is exclusively produced in cells and reflected at the lattice boundary. This is necessary since we do not consider the canalization mode of auxin transport and cannot sustain steady shoot-root flows. Setting auxin inflows and outflows at the boundary would result in artificial depletion and accumulation of auxin at that particular boundary.

In brief, auxin accumulation in the founder cells is aided by an up-the-gradient endodermal auxin flows (Fig. 1 F). This is similar behaviour to that found by (30), where PIN3 transports auxin from the overlying endodermis back to the specified pericycle cells. Because of our model’s high auxin influx carrier density in activated XPP cells, this auxin maximum is stable and persists throughout growth. Auxin flow patterns before and after radial expansion of the activated XPP cells remain stable (Fig. S2).

Since auxin controls growth, we propose cell extensibility, *φ*_cell_, to vary as a function of auxin. The coupling of cell extensibilities (Eq. 6 is rate-limiting, which implies that most growth occurs on the shared interface between activated XPP cells. Yet, there is a challenge with assigning nonzero extensibility to all walls of the simulated tissue. The stress on periclinal walls would artificially decay as growth occurs due to the tissue being confined according to the boundary conditions. To avoid this, we opted for simply having the two activated XPP cells with the same nonzero extensibility (*φ*) and zero everywhere else.

### Auxin accumulation in activated XPP cells leads to modest radial growth and redirects auxin flows towards endodermis

We first examined how an increase in auxin concentration alone contributes to the radial expansion of a pair of founder cells during lateral root initiation. To quantify this expansion, we tracked changes in the length of the shared interface between the two activated XPP cells and their overall morphology. Our simulations showed that growth occurs but saturates around a 13% increase (Fig. 2 A).

**Fig. 2.**
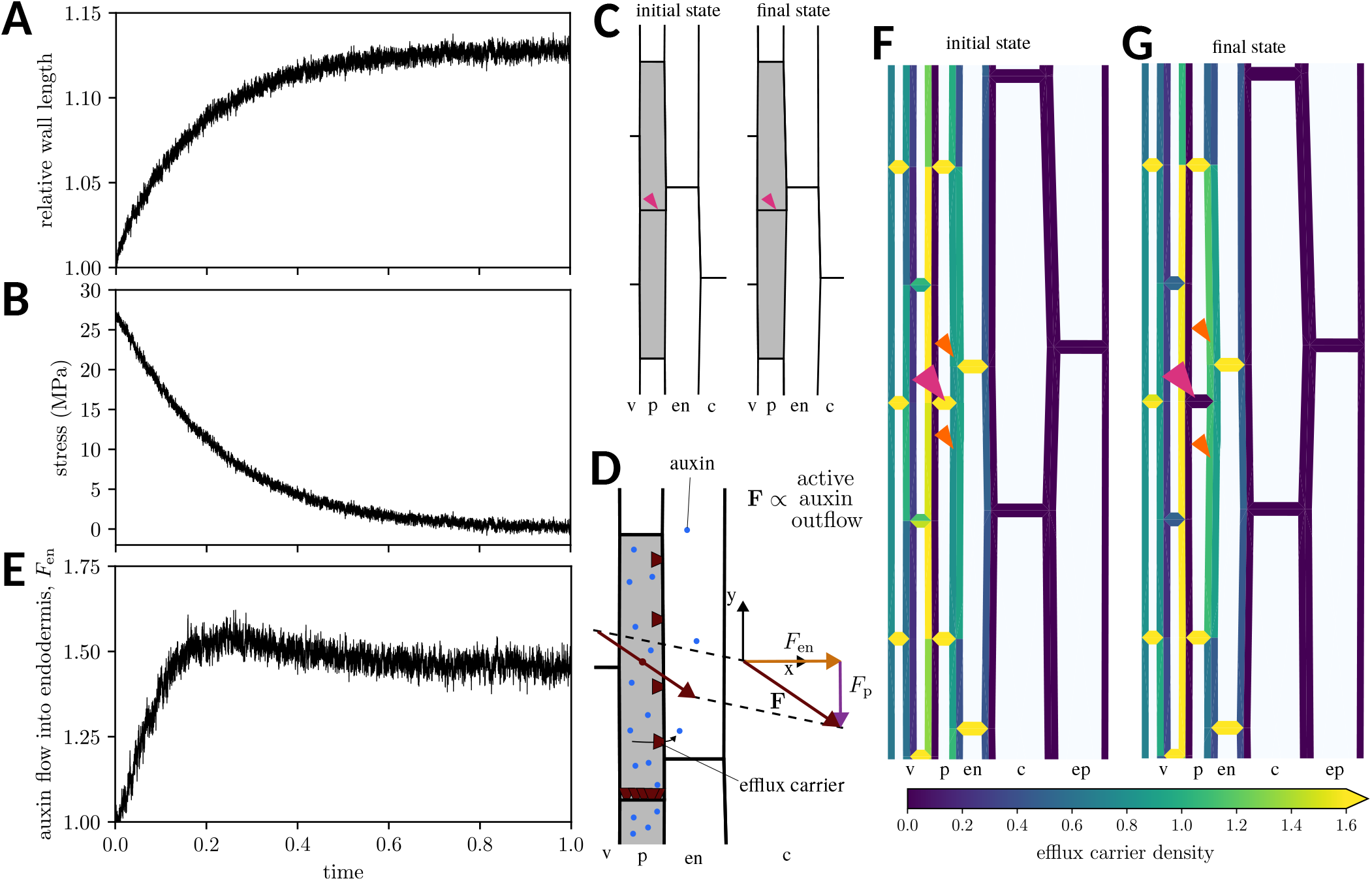
Auxin accumulation in activated XPP cells only triggers modest growth but potentiates auxin flows towards the endodermis. (A) The relative length of the shared wall (pink arrowheads in C) between two activated XPP cells (grey cells) increases over time, reaching a saturation value of around 13%. Accompanying this growth, stress decreases over time (B). The final state of the tissue remains geometrically close to the initial one (C). (D) Computation of the auxin flow. Integrating the density of auxin efflux carriers along the length of the activated XPP cells determines the flow of auxin leaving the cell (**Φ**). This vector can be decomposed in one component toward the endodermis (*F*_en_) and one parallel to the pericycle layer (*F*_p_). (E) The relative intensity of auxin flow leaving the activated XPP cell toward the endodermis (*F*_en_). Efflux carrier density (F) before growth takes place and (G) after growth takes place. Efflux carrier density decreases on the shared wall (pink arrowhead) between activated pericycle cells (shaded cells), and efflux carrier density on the wall facing the endodermis increases (orange arrowheads). In (A), (B) and (E), the measures are normalised to the value at *t* = 0 (initial state). *t* = 1 marks the final state. The different layers are labelled: vasculature (v), pericycle (p), endodermis (en), cortex (c).

This growth coincides with reduced stress within the common wall (Fig. 2 B). However, the overall cell shape remains distinct from the trapezoidal configuration characteristic of activated XPP cells that expand and prepare for asymmetric division (Fig. 2 C).

As stress diminishes within the shared wall connecting the two activated XPP cells, the density of auxin efflux carriers binding to this interface decreases. In our model, carriers compete for binding among different interfaces, leading to the migration of carriers to the following most stressed interface. Initially, the highest density of auxin efflux carriers is in the anticlinal wall between the adjacent activated XPP cells (Fig. 2 F). With growth and stress reduction, carrier density increases at competing cell walls, particularly binding toward the endodermis (Fig. 2 G). Notably, even though the efflux carrier density associated with the endodermis remains relatively low, the cross-sectional area through which auxin flows must be considered. We introduce a measure similar to the polarity concept in (20) to quantify auxin flow due to auxin efflux carriers, showing how the component of auxin flow directed toward the endodermis evolves (Fig. 2 D, E). This measure is directly proportional to the net outflow of auxin (Supplemental Eq. 13). With auxin-induced stress relaxation between adjacent cells, a 40-50% increase in auxin flow exiting the cell toward the endodermis is predicted.

Here, we hypothesize a mechanics-based regulation of auxin transport mechanism, akin to observations in shoot ablation experiments (18, 19, 54). Although the direct link between auxin transport and mechanical stress in roots remains unproven, our model aligns with empirical findings, indicating the importance of theoretical exploration in bridging gaps left by limited experimental data. It suggests the possibility of stress-induced PIN accumulation in roots, not just shoots. While the causal connection is not established (18), there is an observed correlation in shoots between re-orientation of cortical microtubules (CMTs), stress patterns, and auxin efflux carriers post-cell ablation. Similarly, root cell ablation prompts microtubule reorientation around the wound above primed XPP cells (**?**). Directly assessing the correlation between stress pattern changes and auxin efflux carrier polarization post-cell ablation in roots would be an important step to validate the assumptions proposed here.

It’s important to note that our findings on auxin redirection depend on the selected periclinal stress value. Since efflux carriers prefer binding to anticlinal walls (25), we chose a lower periclinal wall stress value, approximately *σ* = 20 MPa. This choice was made to align the initial rest lengths of periclinal walls with the desired overall stress value of a periclinal wall. Selecting lower stress values would obstruct efflux-mediated auxin transport between different layers, whereas higher values would yield an efflux carrier density different from the one empirically observed (Fig. 1 I, J, S2). We only modelled LR initiation until the first round of cell divisions. Given the dynamics of auxin transport patterns past the first anticlinal divisions (30), it would be interesting to investigate how cell division influences growth and auxin patterns.

In summary, our results indicate that increased auxin concentration within adjacent pericycle cells can induce softening of the shared anticlinal wall, resulting in a modest expansion that alone falls short of reproducing the typical cell aspect changes observed *in vivo*. Nevertheless, our model predicts that this stress-induced alteration in auxin efflux carriers subsequently triggers a redistribution, leading to a 40-50% enhancement in auxin flow from the founder cells toward the endodermis.

### Auxin signaling in the endodermis predates the activation of SHY2 expression and the pronounced radial expansion of the activated pericycle

Increasing the auxin flow toward the endodermis may trigger an auxin response in this layer. If this were the case, we predict that auxin responses should first occur in activated XPP cells and then in the endodermis (31). *SHORT HYPOCOTYL 2* (*SHY2*) encodes an AUX/IAA involved in endodermal auxin perception and subsequent endodermal remodelling (31). Controlled transcriptionally by auxin (55), a *SHY2* reporter shows endodermal expression from Stage I onwards (31). To empirically verify that endodermal response to auxin, followed by activation of *SHY2*, is happening *after* redirection of auxin flow from the activated XPP cells, we live-imaged the expression of the nuclear auxin signalling output *DR5v2::Venus* (56) and the *SHY2* transcriptional reporter (31).

Before pericycle activation, neither *SHY2* nor *DR5v2* expression was observed in pericycle or endodermis. Upon activation, *DR5v2* signal appeared in pericycle and contiguous endodermal layers, while *SHY2* expression was solely in pericycle (Fig. 3 A). At Stage I, coinciding with noticeable pericycle swelling (Fig. 3 B), *SHY2* and *DR5v2* were observed in both pericycle and endodermis. This sequential auxin signalling in the endodermis, followed by *SHY2* activation, is in agreement with previous observations (30) and supports the model’s prediction that reversed auxin flow due to modest pericycle expansion can initiate endodermal auxin responses (31).

**Fig. 3.**
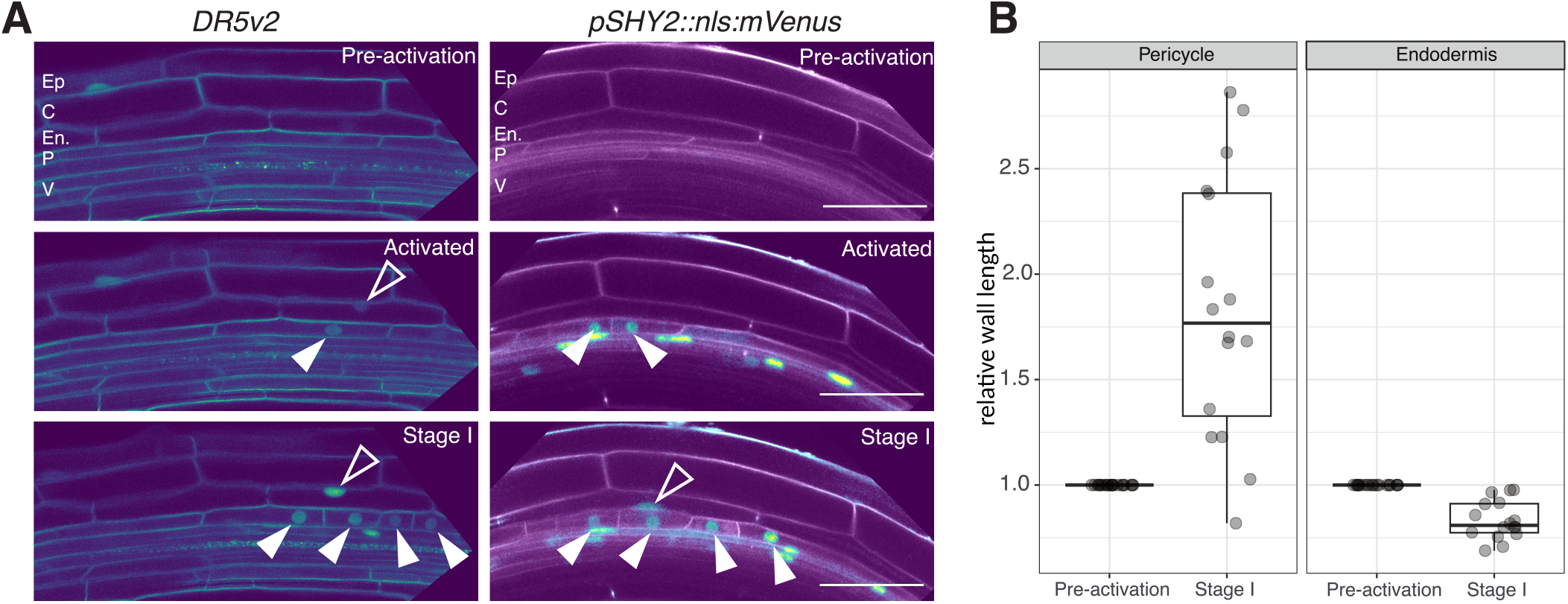
Auxin signaling predates expression of SHY2 in the endodermis and endodermal accommodation. (A) Confocal sections of time-lapse recording monitoring the expression of auxin signalling (*DR5v2*) and *SHY2* expression (*pSHY* 2 :: *nls* : *mV enus*) before pericycle activation, in founder cells and Stage I primordia. The filled arrowheads indicate a signal in the pericycle nuclei, and the open arrowheads signal in the endodermis. Note the *DR5v2* signal in the endodermis above the founder cells before the expression of *SHY2* in this tissue. The different layers are labelled vasculature (V), pericycle (P), endodermis (En), cortex (C) and epidermis (Ep). Of note, the cell outlines are labelled by the plasma membrane marker PIP4;1-GFP (for the *DR5v2* line) and by Propidium iodide (for the *pSHY2::nls:mVenus* line). Images are representative of three biological replicates. Scale bars are 50 *µ*m. (B) Relative pericycle and endodermis cell width (measured at the junction between the two abutting pericycle cells) before activation and at Stage I. Values are for 16 measurements from five different roots before and after pericycle activation.

Our model predicts that auxin accumulation within founder cells induces stress relaxation in the shared anticlinal wall, redirecting auxin flow toward the endodermis. This localized auxin accumulation triggers an auxin signalling response in the endodermis, unleashing the active accommodation of the endodermis to the outgrowing LR. Essential in this process is the mechanical tension in the periclinal interface between the founder cells and the endodermis, which is crucial in coordinating growth and auxin distribution. Root curvature promotes the initiation of LR (57, 58). Curvature in this setting would also increase tension on the periclinal interface between founder cells and endodermis on the convex side of the root (59), which may help auxin-mediated communication between founder cells and the overlying endodermis and further promote auxin flows toward the endodermis upon stress relaxation of the founder cell shared wall.

### Bending modulus of the endodermis limits the radial expansion of the pericycle

Next, we investigated the mechanical factors influencing the expansion of activated XPP cells during lateral root initiation, considering the modest radial expansion induced by initial auxin-driven stress relaxation. We systematically analyzed the impact of various parameters on cell expansion, including the base elastic modulus (*E*_0_) of founder cells and overlying endodermis cells, bending modulus (*k*_*b*_) of overlying endodermal walls (while keeping primed XPP cells at *k*_*b*_ = 100 MPa), as well as turgor pressures in the founder cells (*T*_p_) and the overlying endodermis (*T*_en_) (Fig. S3). The expansion was quantified by measuring the length of the anticlinal wall between the activated XPP cells. Notably, even though we do not explicitly consider water transport across cells, it would result in turgor differences between cells that we explicitly tested here.

We found that reducing the bending modulus (*k*_*b*_) from 3600 MPa to 100 MPa led to non-linear radial expansion, from 13% to 60%, resembling the trapezoidal morphology observed *in planta* (21, 23) (Fig. 4 A, E, F, S3). An increase in pericycle turgor pressure (*T*_p_), especially at very high values (2 MPa), amplified the expansion of the activated XPP cell (Fig. 4 D, G. S3). However, reductions in endodermal turgor (*T*_en_) or elastic modulus (*E*_0_) minimally affected the pericycle expansion (Fig. 4 B, C).

**Fig. 4.**
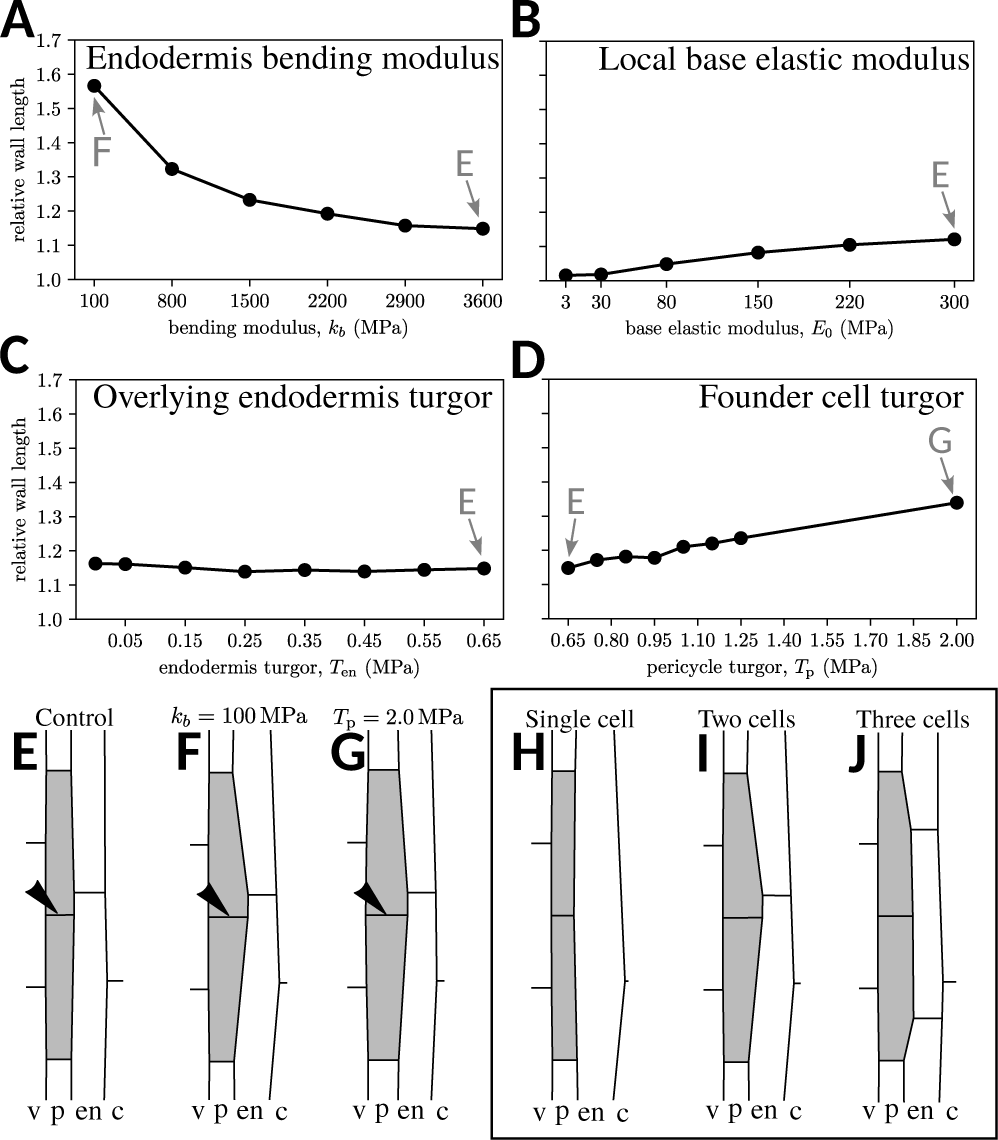
The endodermis’s mechanical properties, topology, and geometry potentiate radial expansion and define LRP shape. (A-D) Relative length of the shared wall between primed XPP cells (grey) for different values of the overlying endodermis and founder cell bending modulus (*k*_*b*_, A), base elastic modulus (*E*_0_, B), overlying endodermis turgor (*T*_en_, C) and founder cell turgor (*T*_p_, D). (E-G) Morphology of the ends-state of the simulations. The grey arrows and letters in (A-D) indicate the corresponding morphologies in (E-G). (H-J) Growth simulation results with one (H), two (I), and three (J) endodermal cells overlying the activated XPP cells. The arrowheads indicate the shared wall between primed XPP cells. The labelled layers are vasculature (v), pericycle (p), endodermis (en), and cortex (c).

These findings suggest that increased pericycle turgor and decreased endodermal resistance to deformation can trigger radial expansion. One notable difference between pericycleturgor-driven and bending-modulus-driven radial expansion lies in the final shape of the founder cells. Upon turgor-driven expansion, the founder cells expanded less toward the endodermis, resulting in a flatter emerging lateral root primordium (LRP) tip. Combining higher pericycle turgor with lower endodermal turgor might rectify this directional bias deficiency. Auxin promotes the expression and activity of pumps and channels that contribute to water influx (60), which could contribute to a local increase of turgor pressure in the pericycle. However, this aspect must be mitigated by the quick water exchange that could equilibrate the turgor pressure since the pericycle cells are still symplastically connected at this point (61). A critical difference is that this shape change is gradual when increasing pericycle turgor and more abrupt when decreasing bending modulus (Fig. 4 A, D, S4). The results suggest that this process is a tug-of-war between turgor and bending modulus, where either turgor increases enough to overcome the bending modulus or bending modulus drops, allowing the existing turgor to drive further radial expansion. Regardless, we have shown radial expansion to be possible with mechanical modulation alone.

The non-linear behaviour of the bending modulus helps explain sudden behaviour changes during this process, whereas the pericycle turgor-driven expansion lacks that particular property on its own. What molecular process could modulate this bending resistance in vivo? In a recent study, we highlighted the distinct organization of cortical microtubules (CMTs) in the endodermis, with an ordered longitudinally oriented array on the pericycle-facing side (34). LR initiation induces reorganization of these microtubules. As CMTs influence wall mechanical properties through cellulose fibril deposition orientation control, the organisation and/or dynamics of the CMTs arrays could thus be a modulator of the bending resistance of the endodermis. Interestingly, we showed that the auxin-dependent induction of the microtubule-associated protein MAP70-5 is necessary to remodel the CMTs (34). Thus MAP70-5 expression could alter CMTs and endodermal wall properties, enhancing founder cell expansion. Supporting this view, abolishing the bending resistance altogether (setting (*k*_*b*_) to 0 MPa) leads to a froth-like behaviour of the tissue (Fig. S5), reminiscent of the aspect of roots cells when treated with the CMT depolymerising agent oryzalin (62). The radial expansion of the shared anticlinal wall primarily occurs after the first anticlinal divisions of founder cells. As our model does not implement cell divisions, investigating how new anticlinal cell interfaces contribute to the radial expansion and how carrier density changes upon cell division could provide valuable insights.

### Endodermis architecture defines LRP shape

We explored whether the arrangement of endodermis cells above the founder cells impacted pericycle growth and shape. Our simulations have mainly focused on lattices where two adjacent endodermal cells overlay the two activated XPP cells, resulting in one endodermis-endodermis interface above the pericycle. We simulated additional cases with either no endodermis-endodermis interface or two such interfaces overlying the pericycle. Employing the same parameters as before (bending modulus of both the activated pericycle cells and overlying endodermal cells at *k*_*b*_ = 100MPa), we present the outcomes (Fig. 4 H-J).

When only one endodermal cell overlays the pericycle, no radial expansion was observed, as there was no strain on the endodermis-pericycle wall (Fig. 4 H). The reduced strain and stress on the wall between the activated XPP cells corresponded to a slower growth rate (Fig. S6 A). To rule out limited degrees of freedom as a causal factor, we conducted simulations where the endodermis-endodermis junction was removed while retaining the respective vertices for displacement. This also resulted in a lack of pericycle expansion (Fig. S6 B, C).

In contrast, the configuration with three overlying endodermal cells led to distinct results (Fig. 4 J). In this scenario, the founder cells expanded to a maximum width, similar to the earlier two-cell configuration (Fig. 4 I, J, Fig. S6 A), albeit while exhibiting slower growth. However, the resulting shape showed noticeable divergence, with the three-cell arrangement producing a flatter tip than the two-cell counterpart.

These findings suggest that the spatial arrangement of endodermis cells governs the shape of the developing primordium, and not all configurations equally support the radial expansion of the primed pericycle. A prior study tracking cell lineage in lateral root primordia consistently observed that the first activated XPP cell to divide was always beneath an endodermal junction (63). Since radial expansion precedes activated XPP cell division (33), this observation is in agreement with our model predictions and underscores the critical role of endodermis topology in channelling the initiation of the lateral root. The dependence of lateral root primordium shape on endodermis topology complicates our understanding of LR initiation. The number and arrangement of overlying endodermal cells dictate the extent and direction of pericycle expansion. As additional founder cells are recruited laterally (32), the implications for LR morphogenesis and tissue morphology modulation during growth are intriguing, especially in a three-dimensional context not captured by our two-dimensional model.

## Conclusion

Our study combines computational modelling and empirical observations to explore the mechanical bases governing the radial expansion of founder cells and endodermal accommodation during lateral root initiation. The central role of auxin in mediating growth and its subsequent impact on mechanical stress distribution emerge as critical components of this process. We find that the mechanical properties of the endodermis are pivotal in shaping the radial expansion of founder cells. Reducing the bending modulus of endodermal walls significantly promotes pericycle growth, highlighting the mechanical regulation of cell expansion. This complements existing knowledge about the role of turgor pressure in growth and emphasizes the importance of localized modifications in cell wall properties for cell growth

## AUTHOR CONTRIBUTIONS

Conceptualization: J. Ramos, K. Alim, A. Maizel; Methodology: J. Ramos, K. Alim; Formal analysis: J. Ramos, K. Alim; Investigation: J. Ramos, B. Reyes-Hernández; Data curation: J. Ramos, B. Reyes-Hernández; Writing-original draft: J. Ramos, A. Maizel; Writing-review & editing: J. Ramos, K. Alim, A. Maizel; Visualization: J. Ramos, B. Reyes-Hernández; Supervision: K. Alim, A. Maizel; Funding acquisition: K. Alim, A. Maizel

## ACKNOWLEDGEMENTS

This work was supported by the German Research Foundation (DFG) through the FOR2581 grants (MA5293/6-2, AL1429/4-1 and AL1429/3-1) and the Consejo Nacional de Ciencia y Tecnología of Mexico (CONACYT) to BJRH (grant 740701). We thank J. Dubrovsky and J. Vermeer for their critical comments on the manuscript.

## SUPPORTING INFORMATION

Supplementary information is available online at: XXXXXX

## PEER REVIEW HISTORY

The peer review history is available online at: XXXXXX

## Bibliography

1. Christopher D. Whitewoods, Beatriz Gonçalves, Jie Cheng, Minlong Cui, Richard Kennaway, Karen Lee, Claire Bushell, Man Yu, Chunlan Piao, and Enrico Coen. Evolution of carnivorous traps from planar leaves through simple shifts in gene expression. Science, 367(6473):91–96, 2020. doi: 10.1126/science.aay5433.

2. Takahiro Yamaguchi, Hirokazu Tsukaya, Hironori Fujita, Kenji Fukushima, Mitsuyasu Hasebe, and Masayoshi Kawaguchi. Oriented cell division shapes carnivorous pitcher leaves of Sarracenia purpurea. Nature Communications, 6(1):1–10, 2015. doi: 10.1038/ncomms7450.

3. Zhaojun Ding and Jiří Friml. Auxin regulates distal stem cell differentiation in Arabidopsis roots. Proceedings of the National Academy of Sciences, 107(26):12046–12051, 2010. doi: 10.1073/pnas.1000672107.

4. Ottoline Leyser. Auxin Signaling. Plant Physiology, 176(1):465–479, jan 2018. ISSN 0032-0889. doi: 10.1104/pp.17.00765.

5. William D. Teale, Ivan A. Paponov, and Klaus Palme. Auxin in action: signalling, transport and the control of plant growth and development. Nature Reviews Molecular Cell Biology, 7 (11):847–859, nov 2006. ISSN 1471-0072. doi: 10.1038/nrm2020.

6. Jodi L Stewart and Jennifer L Nemhauser. Do Trees Grow on Money? Auxin as the Currency of the Cellular Economy. Cold Spring Harbor Perspectives in Biology, 2(2):a001420–a001420, feb 2010. ISSN 1943-0264. doi: 10.1101/cshperspect.a001420.

7. Didier Reinhardt, Eva-Rachele Pesce, Pia Stieger, Therese Mandel, Kurt Baltensperger, Malcolm Bennett, Jan Traas, Jiří Friml, and Cris Kuhlemeier. Regulation of phyllotaxis by polar auxin transport. Nature, 426(6964):255–60, 2003. ISSN 1476-4687. doi: 10.1038/nature02081.

8. Teva Vernoux, Fabrice Besnard, and Jan Traas. Auxin at the Shoot Apical Meristem. Cold Spring Harbor Perspectives in Biology, 2(4):a001487–a001487, apr 2010. ISSN 1943-0264. doi: 10.1101/cshperspect.a001487.

9. Massimiliano Sassi and Teva Vernoux. Auxin and self-organization at the shoot apical meristem. Journal of Experimental Botany, 64(9):2579–2592, jun 2013. ISSN 1460-2431. doi: 10.1093/jxb/ert101.

10. Siobhan A. Braybrook and Alexis Peaucelle. Mechano-Chemical Aspects of Organ Formation in Arabidopsis thaliana: The Relationship between Auxin and Pectin. PLoS ONE, 8(3): e57813, mar 2013. ISSN 1932-6203. doi: 10.1371/journal.pone.0057813.

11. Alexis Peaucelle, Siobhan A. Braybrook, Laurent Le Guillou, Emeric Bron, Cris Kuhlemeier, and Herman Höfte. Pectin-Induced Changes in Cell Wall Mechanics Underlie Organ Initiation in Arabidopsis. Current Biology, 21(20):1720–1726, oct 2011. ISSN 09609822. doi: 10.1016/j.cub.2011.08.057.

12. J. Petrasek and J. Friml. Auxin transport routes in plant development. Development, 136 (16):2675–2688, aug 2009. ISSN 0950-1991. doi: 10.1242/dev.030353.

13. Eric M. Kramer and Malcolm J. Bennett. Auxin transport: a field in flux. Trends in Plant Science, 11(8):382–386, aug 2006. ISSN 13601385. doi: 10.1016/j.tplants.2006.06.002.

14. Richard S Smith. The Role of Auxin Transport in Plant Patterning Mechanisms. PLoS Biology, 6(12):e323, ec 2008. ISSN 1545-7885. doi: 10.1371/journal.pbio.0060323.

15. Didier Reinhardt, Therese Mandel, and Cris Kuhlemeier. Auxin Regulates the Initiation and Radial Position of Plant Lateral Organs. The Plant Cell, 12(4):507, apr 2000. ISSN 10404651. doi: 10.2307/3871065.

16. Richard S Smith, S. Guyomarc’h, Therese Mandel, Didier Reinhardt, Cris Kuhlemeier, and Przemyslaw Prusinkiewicz. A plausible model of phyllotaxis. Proceedings of the National Academy of Sciences USA, 103(5):1301–1306, jan 2006. ISSN 0027-8424. doi: 10.1073/pnas.0510457103.

17. Neha Bhatia, Behruz Bozorg, André Larsson, Carolyn Ohno, Henrik Jönsson, and Marcus G. Heisler. Auxin Acts through MONOPTEROS to Regulate Plant Cell Polarity and Pattern Phyllotaxis. Current Biology, 26(23):3202–3208, ec 2016. ISSN 09609822. doi: 10.1016/j.cub.2016.09.044.

18. Marcus G. Heisler, Olivier Hamant, Pawel Krupinski, Magalie Uyttewaal, Carolyn Ohno, Henrik Jönsson, Jan Traas, and Elliot M. Meyerowitz. Alignment between PIN1 Polarity and Microtubule Orientation in the Shoot Apical Meristem Reveals a Tight Coupling between Morphogenesis and Auxin Transport. PLoS Biology, 8(10):e1000516, oct 2010. ISSN 1545-7885. doi: 10.1371/journal.pbio.1000516.

19. Naomi Nakayama, Richard S Smith, Therese Mandel, Sarah Robinson, Seisuke Kimura, Arezki Boudaoud, and Cris Kuhlemeier. Mechanical regulation of auxin-mediated growth. Current Biology, 22(16):1468–1476, August 2012.

20. D. Ramos, João R., Maizel, Alexis, and Alim, Karen. Tissue-wide integration of mechanical cues promotes effective auxin patterning. Eur. Phys. J. Plus, 136(2):250, 2021. doi: 10.1140/epjp/s13360-021-01204-6.

21. Amaya Vilches-Barro and Alexis Maizel. Talking through walls: Mechanisms of lateral root emergence in Arabidopsis thaliana. Current Opinion in Plant Biology, 23(May):31–38, 2015. ISSN 13695266. doi: 10.1016/j.pbi.2014.10.005.

22. Dorothee Stoeckle, Martha Thellmann, and Joop EM Vermeer. Breakout — lateral root emergence in Arabidopsis thaliana. Current Opinion in Plant Biology, 41:67–72, feb 2018. ISSN 13695266. doi: 10.1016/j.pbi.2017.09.005.

23. Yujuan Du and Ben Scheres. Lateral root formation and the multiple roles of auxin. Journal of Experimental Botany, 69(2):155–167, 07 2017. ISSN 0022-0957. doi: 10.1093/jxb/erx223.

24. Joseph G. Dubrovsky, Michael Sauer, Selene Napsucialy-Mendivil, Maria G. Ivanchenko, Jiří Friml, Svetlana Shishkova, John Celenza, and E. Benkova. Auxin acts as a local morphogenetic trigger to specify lateral root founder cells. Proceedings of the National Academy of Sciences USA, 105(25):8790–8794, jun 2008. ISSN 0027-8424. doi: 10.1073/pnas.0712307105.

25. Eva Benková, Marta Michniewicz, Michael Sauer, Thomas Teichmann, Daniela Seifertová, Gerd Jürgens, and Jiřı Friml. Local, Efflux-Dependent Auxin Gradients as a Common Module for Plant Organ Formation. Cell, 115(5):591–602, nov 2003. ISSN 00928674. doi: 10.1016/S0092-8674(03)00924-3.

26. Anna N. Stepanova, Joyce Robertson-Hoyt, Jeonga Yun, Larissa M. Benavente, De-Yu Xie, Karel Doležal, Alexandra Schlereth, Gerd Jürgens, and Jose M. Alonso. TAA1-mediated auxin biosynthesis is essential for hormone crosstalk and plant development. Cell, 133(1): 177–191, Apr 2008. ISSN 0092-8674. doi: 10.1016/j.cell.2008.01.047.

27. Alan Marchant, Rishikesh Bhalerao, Ilda Casimiro, Jan Eklöf, Pedro J. Casero, Malcolm Bennett, and Goran Sandberg. AUX1 Promotes Lateral Root Formation by Facilitating Indole-3-Acetic Acid Distribution between Sink and Source Tissues in the Arabidopsis Seedling. The Plant Cell, 14(3):589–597, mar 2002. ISSN 1040-4651. doi: 10.1105/tpc.010354.

28. Kamal Swarup, Eva Benková, Ranjan Swarup, Ilda Casimiro, Benjamin Péret, Yaodong Yang, Geraint Parry, Erik Nielsen, Ive De Smet, Steffen Vanneste, Mitch P. Levesque, David Carrier, Nicholas James, Vanessa Calvo, Karin Ljung, Eric Kramer, Rebecca Roberts, Neil Graham, Sylvestre Marillonnet, Kanu Patel, Jonathan D.G. Jones, Christopher G. Taylor, Daniel P. Schachtman, Sean May, Goran Sandberg, Philip Benfey, Jiri Friml, Ian Kerr, Tom Beeckman, Laurent Laplaze, and Malcolm J. Bennett. The auxin influx carrier LAX3 promotes lateral root emergence. Nature Cell Biology, 10(8):946–954, aug 2008. ISSN 1465-7392. doi: 10.1038/ncb1754.

29. B. Péret, A. M. Middleton, A. P. French, A. Larrieu, A. Bishopp, M. Njo, D. M. Wells, S. Porco, N. Mellor, L. R. Band, I. Casimiro, J. Kleine-Vehn, S. Vanneste, I. Sairanen, R. Mallet, G. Sandberg, K. Ljung, T. Beeckman, E. Benková, J. Friml, E. Kramer, J. R. King, I. De Smet, T. Pridmore, M. Owen, and M. J. Bennett. Sequential induction of auxin efflux and influx carriers regulates lateral root emergence. Molecular Systems Biology, 9(1):699–699, apr 2014. ISSN 1744-4292. doi: 10.1038/msb.2013.43.

30. Peter Marhavý, Marleen Vanstraelen, Bert De Rybel, Ding Zhaojun, Malcolm J. Bennett, Tom Beeckman, and Eva Benková. Auxin reflux between the endodermis and pericycle promotes lateral root initiation. The EMBO Journal, 32(1):149–158, nov 2012. ISSN 0261-4189. doi: 10.1038/emboj.2012.303.

31. J. E. M. Vermeer, D. von Wangenheim, M. Barberon, Y. Lee, E. H. K. Stelzer, A. Maizel, and N. Geldner. A Spatial Accommodation by Neighboring Cells Is Required for Organ Initiation in Arabidopsis. Science, 343(6167):178–183, jan 2014. ISSN 0036-8075. doi: 10.1126/science.1245871.

32. Héctor H Torres-Martínez, Paul Hernández-Herrera, Gabriel Corkidi, and Joseph G Dubrovsky. From one cell to many: Morphogenetic field of lateral root founder cells in arabidopsis thaliana is built by gradual recruitment. Proceedings of the National Academy of Sciences, 117(34):20943–20949, August 2020.

33. Amaya Vilches Barro, Dorothee Stöckle, Martha Thellmann, Paola Ruiz-Duarte, Lotte Bald, Marion Louveaux, Patrick von Born, Philipp Denninger, Tatsuaki Goh, Hidehiro Fukaki, Joop E.M. Vermeer, and Alexis Maizel. Cytoskeleton dynamics are necessary for early events of lateral root initiation in arabidopsis. Current Biology, 29(15):2443–2454.e5, August 2019. doi: 10.1016/j.cub.2019.06.039.

34. Dorothee Stöckle, Blanca Jazmin Reyes-Hernández, Amaya Vilches Barro, Milica Nenadić, Zsofiá Winter, Sophie Marc-Martin, Lotte Bald, Robertas Ursache, Satoshi Fujita, Alexis Maizel, and Joop EM Vermeer. Microtubule-based perception of mechanical conflicts controls plant organ morphogenesis. Science Advances, 8(6):eabm4974, 2022. doi: 10.1126/sciadv.abm4974.

35. Jean-Daniel Julien, Alain Pumir, and Arezki Boudaoud. Strain-or stress-sensing in mechanochemical patterning by the phytohormone auxin. bioRxiv, 2019. doi: 10.1101/582551.

36. Roeland M.H. Merks, Michael Guravage, Dirk Inzé, and Gerrit T.S. Beemster. Virtualleaf: An open-source framework for cell-based modeling of plant tissue growth and development. Plant Physiology, 155(2):656–666, 02 2011. ISSN 0032-0889. doi: 10.1104/pp.110.167619.

37. D. B. Staple, R. Farhadifar, J. C. Röper, B. Aigouy, S. Eaton, and F. Jülicher. Mechanics and remodelling of cell packings in epithelia. The European Physical Journal E, 33(2):117–127, oct 2010. ISSN 1292-8941. doi: 10.1140/epje/i2010-10677-0.

38. Silvanus Alt, Poulami Ganguly, and Guillaume Salbreux. Vertex models: from cell mechanics to tissue morphogenesis. Philosophical Transactions of the Royal Society B: Biological Sciences, 372(1720):20150520, may 2017. ISSN 0962-8436. doi: 10.1098/rstb.2015.0520.

39. Jason Khadka, Jean-Daniel Julien, and Karen Alim. Feedback from tissue mechanics self-organizes efficient outgrowth of plant organ. Biophysical Journal, 117(10):1995–2004, nov 2019. doi: 10.1016/j.bpj.2019.10.019.

40. H. Jonsson, M. G. Heisler, B. E. Shapiro, E. M. Meyerowitz, and E. Mjolsness. An auxin-driven polarized transport model for phyllotaxis. Proceedings of the National Academy of Sciences USA, 103(5):1633–1638, jan 2006. ISSN 0027-8424. doi: 10.1073/pnas.0509839103.

41. Patrik Sahlin B. Söderberg, and Henrik Jönsson. Regulated transport as a mechanism for pattern generation: Capabilities for phyllotaxis and beyond. Journal of Theoretical Biology, 258(1):60–70, may 2009. ISSN 00225193. doi: 10.1016/j.jtbi.2009.01.019.

42. F Corson, O Hamant, S Bohn, J Traas, A Boudaoud, and Y Couder. Turning a plant tissue into a living cell froth through isotropic growth. Proceedings of the National Academy of Sciences USA, 106(21):8453–8458, may 2009. ISSN 0027-8424. doi: 10.1073/pnas.0812493106.

43. Yuchen Long, Ibrahim Cheddadi, Gabriella Mosca, Vincent Mirabet, Mathilde Dumond, Annamaria Kiss, Jan Traas, Christophe Godin, and Arezki Boudaoud. Cellular heterogeneity in pressure and growth emerges from tissue topology and geometry. Current Biology, 30 (8):1504–1516.e8, 2020. ISSN 0960-9822. doi: 10.1016/j.cub.2020.02.027.

44. L. Livadaru, R. R. Netz, and H. J. Kreuzer. Stretching response of discrete semiflexible polymers. Macromolecules, 36(10):3732–3744, April 2003. doi: 10.1021/ma020751g.

45. J. Kierfeld, O. Niamploy, V. Sa-yakanit, and R. Lipowsky. Stretching of semiflexible polymers with elastic bonds. The European Physical Journal E, 14(1):17–34, May 2004. doi: 10.1140/epje/i2003-10089-3.

46. Jan Kierfeld, Krzysztof Baczynski, Petra Gutjahr, and Reinhard Lipowsky. Semiflexible polymers and filaments: From variational problems to fluctuations. AIP Conference Proceedings, 1002:151, 04 2008. doi: 10.1063/1.2918091.

47. James A. Lockhart. An analysis of irreversible plant cell elongation. Journal of Theoretical Biology, 8(2):264–275, mar 1965. ISSN 00225193. doi: 10.1016/0022-5193(65)90077-9.

48. J. K. E. Ortega. Augmented Growth Equation for Cell Wall Expansion. Plant Physiology, 79 (1):318–320, sep 1985. ISSN 0032-0889. doi: 10.1104/pp.79.1.318.

49. P. Heckbert. Quad-edge data structure library.

50. S. G. Johnson. The nlopt nonlinear-optimization package.

51. Charles R. Harris, K. Jarrod Millman, Stéfan J van der Walt, Ralf Gommers, Pauli Virtanen, David Cournapeau, Eric Wieser, Julian Taylor, Sebastian Berg, Nathaniel J. Smith, Robert Kern, Matti Picus, Stephan Hoyer, Marten H. van Kerkwijk, Matthew Brett, Allan Haldane, Jaime Fernández del Río, Mark Wiebe, Pearu Peterson, Pierre Gérard-Marchant, Kevin Sheppard, Tyler Reddy, Warren Weckesser, Hameer Abbasi, Christoph Gohlke, and Travis E. Oliphant. Array programming with NumPy. Nature, 585:357–362, 2020. doi: 10.1038/s41586-020-2649-2.

52. J. D. Hunter. Matplotlib: A 2d graphics environment. Computing in Science & Engineering, 9(3):90–95, 2007. doi: 10.1109/MCSE.2007.55.

53. Peter Marhavý and Eva Benková. Real-time analysis of lateral root organogenesis in arabidopsis. Bio Protoc, 5(8), April 2015.

54. O. Hamant, M. G. Heisler, H. Jönsson, P. Krupinski, M. Uyttewaal, P. Bokov, F. Corson, P. Sahlin, A. Boudaoud, E. M. Meyerowitz, Y. Couder, and J. Traas. Developmental Patterning by Mechanical Signals in Arabidopsis. Science, 322(5908):1650–1655, dec 2008. ISSN 0036-8075. doi: 10.1126/science.1165594.

55. Qing Tian, Nicholas J. Uhlir, and Jason W. Reed. Arabidopsis SHY2/IAA3 Inhibits Auxin-Regulated Gene Expression. The Plant Cell, 14(2):301–319, 02 2002. ISSN 1040-4651. doi: 10.1105/tpc.010283.

56. D.R. Boer, A. Freire Rios, W.A.M. van den Berg, T. Saaki, I.W. Manfield, S. Kepinski, I. López-Vidrieo, J.M. Franco-Zorilla, S.C. de Vries, R. Solano, D. Weijers, and M. Coll. Structural basis for dna binding specificity by the auxin-dependent arf transcription factors. Cell, 156:577–589, 2014. ISSN 0092-8674. doi: 10.1016/j.cell.2013.12.027.

57. Stefan Kircher and Peter Schopfer. Priming and positioning of lateral roots in Arabidopsis. An approach for an integrating concept. Journal of Experimental Botany, 67(5):1411–1420, 12 2015. ISSN 0022-0957. doi: 10.1093/jxb/erv541.

58. Marta Laskowski, Verônica A Grieneisen, Hugo Hofhuis, Colette A. ten Hove, Paulien Hogeweg, Athanasius F. M Marée, and Ben Scheres. Root system architecture from coupling cell shape to auxin transport. PLOS Biology, 6(12):1–15, 12 2008. doi: 10.1371/journal.pbio.0060307.

59. L D Landau, L. P. Pitaevskii, A. M. Kosevich, and E.M. Lifshitz. Theory of Elasticity. 12 2012. ISBN 9780080570693.

60. Katrin Philippar, Natalya Ivashikina, Peter Ache, May Christian, Hartwig Lüthen, Klaus Palme, and Rainer Hedrich. Auxin activates kat1 and kat2, two k+-channel genes expressed in seedlings of arabidopsis thaliana. The Plant Journal, 37(6):815–827, 2004. doi: 10.1111/j.1365-313X.2003.02006.x.

61. Yoselin Benitez-Alfonso, Christine Faulkner, Ali Pendle, Shunsuke Miyashima, Ykä Helariutta, and Andrew Maule. Symplastic Intercellular Connectivity Regulates Lateral Root Patterning. Developmental Cell, 26(2):136–147, jul 2013. ISSN 15345807. doi: 10.1016/j.devcel.2013.06.010.

62. Tobias I. Baskin, Gerrit T.S. Beemster, Jan E. Judy-March, and Francoise Marga. Disorganization of Cortical Microtubules Stimulates Tangential Expansion and Reduces the Uniformity of Cellulose Microfibril Alignment among Cells in the Root of Arabidopsis. Plant Physiology, 135(4):2279–2290, 08 2004. ISSN 0032-0889. doi: 10.1104/pp.104.040493.

63. Daniel von Wangenheim, Jens Fangerau, Alexander Schmitz, Richard S Smith, Heike Leitte, Ernst H K Stelzer, and Alexis Maizel. Rules and Self-Organizing properties of post-embryonic plant organ cell division patterns. Curr. Biol., 26(4):439–449, January 2016.

