## Supplemental Material for "Auxin-mediated stress relaxation in pericycle and endoderm remodelling drive lateral root initiation"

### 1 Detailed model description

We model the lateral root system using a vertex mechanics and growth model coupled with a compartment model for active and passive auxin transport. Vertex models have described many biological systems [1] and have been employed before in describing plant tissue mechanics [12, 13]. The model we use here builds and expands on a similar formalism used in a previous work [7]. This formalism is reintroduced here and differs slightly from the one presented in the main text.

#### 1.1 Conventions

To describe the geometry of the tissue, we concern ourselves with a set of $N$ vertices with positions $\mathbf{x}_1, \mathbf{x}_2, \ldots, \mathbf{x}_N$, and with the set of edges that connect a vertex to another. We use the indices of the vertices this edge connects to refer to an edge. Properties of an edge, for instance, its length $l$, will be written as $l_{i\,j}$, $i$ and $j$ being different vertices. In this example, it does not matter which index is written down first, yet that is not always the case. Some quantities are defined for one of the cells that edge separates. Stress and efflux carrier density are two prominent examples of this throughout this text. Namely, the subscript $ij$ will be specific to the cell on the left when travelling from $i$ to $j$. Furthermore, it is convenient to separate the labelling of cells and vertices. Whereas we opted to employ Greek letters when referring to quantities of a cell; for example, $A_\alpha$ is the area of cell $\alpha$.

A recurring pattern is the summation of all edges or vertices surrounding a particular cell. To encapsulate this, we define a sequence of vertices around a cell with an arbitrary starting point, $\mathcal{V}_\alpha$, such that for $i \in \mathcal{V}_\alpha$, $i+1$ refers to the next counterclockwise vertex around the cell. This sequence is also cyclic such that, for a cell with $N^{\text{vertices}}$ vertices, $i = 1, 2, \ldots, N^{\text{vertices}}$ and $i = N^{\text{vertices}} + 1$ circles back to $i = 1$. For instance, we can write perimeter of cell $\alpha$ as simply

$$L_\alpha = \sum_{i \in \mathcal{V}_\alpha} l_{i\,i+1}, \tag{1}$$

Sum over $i \in \mathcal{V}_\alpha$ meaning sum over all vertices around the cell in a counterclockwise fashion. In this case, the ordering of the subscript does not matter, but it will in general.

#### 1.2 Mechanical model

We express the mechanical energy ($\mathcal{H}^{\text{mech}}$) as the combination of three different contributions: the longitudinal elastic deformation of cell-cell interfaces represented by edges ($\mathcal{H}_\alpha^{\text{elastic}}$), the resistance to bending at junctions exhibited by plant cells ($\mathcal{H}_\alpha^{\text{bending}}$), and the pressure within the cell ($\mathcal{H}_\alpha^{\text{pressure}}$). For $M$ cells, the mechanical energy is given by

$$\mathcal{H}^{\text{mech}} = \sum_{\alpha=1}^{M} \mathcal{H}_\alpha^{\text{mech}}, \qquad \mathcal{H}_\alpha^{\text{mech}} = \mathcal{H}_\alpha^{\text{elastic}} + \mathcal{H}_\alpha^{\text{bending}} + \mathcal{H}_\alpha^{\text{pressure}}. \tag{2}$$

Each time the tissue changes mechanically, we find a new geometrical configuration by minimizing the mechanical energy with respect to vertex coordinates. We assume this is the shape the tissue will relax to after such perturbation. To speed up convergence, the gradient of mechanical energy is extremely useful since this whole process must be repeated each time auxin concentrations change or growth occurs.

In the following, we detail each energy term.

##### 1.2.1 Longitudinal elastic deformation of the cell wall ($\mathcal{H}_\alpha^{\text{elastic}}$)

Our tissue description considers each edge as representing a cell-cell interface. We will call these walls for simplicity. Each cell is characterized by a unique elastic modulus, $E_\alpha$, and the specific elastic modulus of the wall will be a combination of the moduli of the surrounding cells. Namely, by associating elastic elements in parallel with the properties of each cell, we find the elastic modulus of the wall to be the average of the elastic modulus of both surrounding cells. Because the auxin pattern formation mechanism relies on an asymmetric stress perception by neighbouring cells [9, 12, 7], we will, at no loss of generality, separate each wall into two compartments, one facing each cell on each side of the wall.

In keeping with the notation described above, $E_{i\,j} = E_\alpha$ refers to the wall compartment facing cell $\alpha$, which in this case has to be located on the left of the edge $ij$ when travelling from $i$ to $j$ for the equality to hold true. For each wall, we also attribute a parameter representing the length the wall would have if no forces are applied to it, $l_0$. Let $l_{0\,i\,j}$ represent the rest length of a wall compartment. We assume it to be the same value for both compartments adjacent to the same edge, i.e., $l_{0\,i\,j} = l_{0\,j\,i}$.

Extending and compressing walls cost energy and, as such, we write down the elastic energy density for a single compartment as

$$\psi_{i\,j}^{\text{elastic}} = \frac{1}{2} E_{i\,j} \left( \frac{l_{i\,j} - l_{0\,i\,j}}{l_{0\,i\,j}} \right)^2, \tag{3}$$

which, by integrating over the volume of the wall and summing over all walls around a cell, yields

$$\mathcal{H}_\alpha^{\text{elastic}} = \frac{1}{2} h w E_\alpha \sum_{i \in \mathcal{V}_\alpha} l_{0\,i\,i+1} \left( \frac{l_{i\,i+1} - l_{0\,i\,i+1}}{l_{0\,i\,i+1}} \right)^2, \tag{4}$$

where $h$ and $w$ represent the height and the width of a cell wall compartment (consider $w$ to be the width of half the entire cell-cell interface), and by taking advantage of the fact that $E_\alpha = E_{i\,i+1}, \forall i \in \mathcal{V}_\alpha$.

##### 1.2.2 Bending energy of cell junctions ($\mathcal{H}_\alpha^{\text{bending}}$)

Our choice for bending energy surface density follows that of Canham-Helfrich [8] at position $\mathbf{x}_i$ on the surface of cell $\alpha$,

$$\psi_\alpha^{\text{bending}}(\mathbf{x}_i) = \frac{k_{1\,\alpha}}{2} \left( H(\mathbf{x}_i) - H_0 \right)^2 + k_{2\,\alpha} K(\mathbf{x}_i), \tag{5}$$

where $k_1$ and $k_2$ are bending moduli, $H$ is the mean curvature, and $K$ is the Gaussian curvature. Furthermore, mean and Gaussian curvatures are often decomposed into their eigenvalues, $H = (\kappa_1 + \kappa_2)/2$ and $K = \kappa_1 \kappa_2$. Since our model is two-dimensional, the curvature is a scalar, and eigendecomposition is redundant, leaving us to disregard the second term and adopt $H = \kappa$. Because our cells are polygons, the curvature is non-zero only at cell junctions and, explained in detail here [6], is given by

$$\kappa_i \approx \frac{2\theta_i}{l_{i-1\,i} + l_{i\,i+1}}, i \in \mathcal{V}_\alpha \tag{6}$$

where $\theta_i$ is the external angle at vertex $i$ made by edges $i-1\,i$ and $i\,i+1$. One may set $H_0$ as a simple parameter, but for this particular system, it is more convenient to set, not to the curvature it would have at rest, but to the angle it would have at rest, $\theta_0$. Moreover, having only one parameter, we rename $k_1$ to $k_b$, the bending modulus. With these simplifications and assumptions, we can write the energy density for each vertex around $\alpha$,

$$\psi_{\alpha\,i}^{\text{bending}} \approx 2 k_{b\,\alpha} \left( \frac{\theta_i - \theta_{0\,i}}{l_{i-1\,i} + l_{i\,i+1}} \right)^2. \tag{7}$$

All that is left is to discretize the perimeter of the polygon. We choose the Voronoi region around the vertex, i.e., the set of closest points. This means that the integration volume we choose is $(l_{i-1\,i} + l_{i\,i+1})/2$, the wall length up until the halfway point of each adjacent edge. Assuming constant energy density, integrating, and summing over vertices around a polygon yields

$$\mathcal{H}_\alpha^{\text{bending}} \approx 2k_{b_\alpha} \sum_{i \in \mathcal{V}_\alpha} \frac{1 - \cos(\theta_i - \theta_{0\,i})}{l_{i-1\,i} + l_{i\,i+1}}, \tag{8}$$

where we have also made use of the Taylor expansion of $\cos(x - x_0) \approx 1 - (x - x_0)^2/2$. This end expression is similar to those often used in semi-flexible discrete polymer chain mechanics [19, 15, 14].

Our choice of wall discretization into single segments precludes us from describing wall curvature along each wall segment. This is not an issue when adjacent cells have the same turgor pressure since the preferred positioning of walls between cells of the same turgor pressure are straight lines [5]. That said, we lose the positive curvature of a wall segment associated with turgid cells and the shape of the epidermal interface with the outer medium. When we test different values of turgor, we, therefore, lose some information on the exact shape cells will take and have to account for it in our interpretation of the results. This also means the changes we do observe are more likely to be significant.

##### 1.2.3 Pressurizing the cell ($\mathcal{H}_\alpha^{\text{pressure}}$)

Each cell of area, $A_\alpha$, in the tissue exerts pressure on its surrounding cell wall. We describe this behaviour by adding a pressure term into our mechanical energy,

$$\mathcal{H}_\alpha^{\text{pressure}} = -T_\alpha A_\alpha, \tag{9}$$

where $T_\alpha$ represents the turgor pressure cell $\alpha$ is under. This way, the cell will expand in any way it can.

#### 1.3 Cell growth model

The cell walls can extend and grow, and to capture this effect, we employ a Maxwell material model to describe this growth behaviour. As laid out by [20, 5], we evolve the rest lengths, $l_0$, over time as a function of stress, $\sigma$, with the following expression, for a single general element with parameters $E$, $l_0$ and length $l$,

$$\frac{1}{l_0}\frac{dl_0}{dt} = \begin{cases} \varphi E \frac{l - l_0}{l_0}, & \sigma > 0, \\ 0, & \sigma \leq 0, \end{cases} \tag{10}$$

where $\varphi$ is the reciprocal of viscosity, which we will refer to as extensibility. Equivalently, the rheological model for this single element is given by Fig. S1 A. These parameters will be assigned to each cell, $\varphi_\alpha$ being the extensibility of cell $\alpha$, and, if $\alpha$ lies on the left of the edge from $i$ to $j$, $\varphi_\alpha = \varphi_{i\,j}$ in a similar manner to the elastic modulus. In order to accommodate different parameters on different sides of the cell wall, the rheological model of the cell wall we use here is given in Fig. S1 B. We intend to keep $l_0$ the same for both wall compartments to keep their phenomenological meaning, so they should evolve with the same equation. It turns out that the model in Fig. S1 B admits a reduced model as in Fig. S1 A, where we combine the parameters $E$ and $\varphi$ of the two cells surrounding a specific wall. The expression for growth for this equivalent model, with the notation presented in this text, would be written as,

$$\frac{1}{l_{0\,i\,j}}\frac{dl_{0\,i\,j}}{dt} = \bar{\varphi}_{i\,j}\bar{E}_{i\,j}\frac{l_{i\,j} - l_{0\,i\,j}}{l_{0\,i\,j}}, \qquad \bar{E}_{i\,j} = \frac{E_{i\,j} + E_{j\,i}}{2} = \bar{E}_{j\,i}, \qquad \bar{\varphi}_{i\,j} = \frac{2\varphi_{i\,j}\varphi_{j\,i}}{\varphi_{i\,j} + \varphi_{j\,i}} = \bar{\varphi}_{j\,i}, \tag{11}$$

for positive stress, leaving us with $l_{0\,i\,j} = l_{0\,j\,i}$ and $dl_{0\,i\,j}/dt = dl_{0\,j\,i}/dt$, if initial values for $l_0$ are the same. Because of the shared value of $l_0$, rather than being exactly that of two parallel Maxwell materials, the material in question is a combination of two elastic parallel elements with two viscous parallel elements in series, ensuring the reversible and irreversible strains are separable (Fig. S1 B).[1] This coupling is also interesting since it tells us that the cell wall growth is limited by the slowest growing component, implying the need for at least two adjacent cells prone to growth for any change to take effect.

An additional complexity of describing plant growth is that plant cells exhibit a threshold of strain or stress below which they cannot grow and need to overcome [22]. Even though we do not take this into account in this work, one could implement it by introducing a step function with a yield value of stress or strain ($\sigma_Y$ or $\varepsilon_Y$), for example, $\varphi E \max\left(0, \left(l - l_0\right)/l_0 - \varepsilon_{\mathrm{Y}}\right)$, or $\varphi \max\left(0, E\left(l - l_0\right)/l_0 - \sigma_{\mathrm{Y}}\right)$. As an example, a similar approach was taken [21]. If the stress of a wall cannot be sustained, as is our case for the shared wall between primed XPP cells before endodermis accommodation, stress of this wall decays with growth. By adding yield stress as mentioned above, then stress would only decay to a base value of $\sigma_{\mathrm{Y}}$, and this could explain why we still see transporters bound to the shared wall of activated XPP cells after lateral root initiation. Interestingly, we observe from our results (Fig. 2 B, E) that, by the time stress on the shared wall reaches 10 MPa, the relocation of efflux carriers has already taken place, meaning that modelling with a yield stress, $\sigma_Y = 10\,\mathrm{MPa}$, would have little to no impact on the final value of auxin flow into the endodermis. However, it would take longer to reach that value.

Given auxin's role during growth, it is reasonable to model extensibility as a function of auxin. Exploring this interaction with auxin flows in other contexts could be interesting. Of particular note is the shoot apical meristem system, where special attention is required since these epithelial cells are not confined, and stress relaxation can be avoided by expanding in a third dimension.

In our simulation, we keep $\varphi_\alpha$ as constant parameters of the model. This is simply a timescale separation approximation because auxin concentration, and hence extensibility, remains high only in the founder cells due to the high density of auxin importers and low everywhere else. This is because the lowest extensible cell governs the behaviour of growth. Additionally, because in our setup we confine the tissue along the root-shoot axis (Fig. 1), slow growth of periclinal walls would cause stress to decay artificially, where *in vivo* this piece of tissue would expand vertically and sustained turgor would lead to sustained stress. Another way to interpret this approximation is to tie extensibility not to auxin concentration itself but to auxin response factor shown to be exclusive to founder cells before activating the accommodation response in the endodermis [3] (and Fig. 3 of the main text).

#### 1.4 Auxin transport model and coupling to mechanics

Our description of auxin transport is similar to the one in [7], which is based on previous approaches by [9, 11, 26].

##### 1.4.1 Auxin transport model

On top of the above mechanical description, we place an auxin transport model between neighbouring cells. Each cell is given an auxin concentration, $a_\alpha$, which is subject to being produced, decaying, and passively and actively transported. Following [7] (namely, the derivation in the SI of this article), we

[1]Consider two such compartments, A and B. Under instantaneous deformations, the elastic response is $\sigma = \left(\sigma_A + \sigma_B\right)/2 = \left(E_A + E_B\right)\varepsilon/2 = \bar{E}\varepsilon$, since the same strain applies to the two parallel branches, and where the factor of 2 accounts for the cross-section difference from considering half a wall to an equivalent wall of combined cross-sections. In contrast, the parallel association of viscous elements will have the same strain rate $\dot{\varepsilon} = \sigma_A\varphi_A = \sigma_B\varphi_B$, since we have forced the same $l_0$. An equivalent Maxwell material would behave as $\dot{\varepsilon} = \sigma\bar{\varphi}$. One can also rewrite $\sigma = \left(1 + \varphi_A/\varphi_B\right)\sigma_A/2$ and finally compute $\bar{\varphi} = \dot{\varepsilon}/\sigma = 2\varphi_A/\left(1 + \varphi_A/\varphi_B\right) = 2\varphi_A\varphi_B/\left(\varphi_A + \varphi_B\right)$

write down the adimensional version of the model,

$$\begin{aligned}\dot{a}_\alpha = 1 - a_\alpha + D \sum_{i\in\mathcal{V}_\alpha} \frac{l_{i\,i+1}}{A_\alpha}\left(a_{\beta(i\,i+1)} - a_\alpha\right)\\ + P \sum_{i\in\mathcal{V}_\alpha} \frac{l_{i\,i+1}}{A_\alpha}\left(p_{i+1\,i}\frac{a_{\beta(i+1\,i)}}{K_{\text{efflux}} + a_{\beta(i+1\,i)}} - p_{i\,i+1}\frac{a_\alpha}{K_{\text{efflux}} + a_\alpha}\right)\\ + Q \sum_{i\in\mathcal{V}_\alpha} \frac{l_{i\,i+1}}{A_\alpha}\left(q_{i\,i+1}\frac{a_{\beta(i+1\,i)}}{K_{\text{influx}} + a_{\beta(i+1\,i)}} - q_{i+1\,i}\frac{a_\alpha}{K_{\text{influx}} + a_\alpha}\right),\end{aligned} \tag{12}$$

where $D$, $P$, $Q$ denote the diffusion strength, and auxin exporter and importer permeabilities respectively, the function $\beta\,(i\,j)$ returns the cell on the left of the edge $i\,j$ when going from $i$ to $j$, and $K_{\text{efflux}}$ and $K_{\text{influx}}$ are constants that determine how quickly the respective auxin carriers saturate in activity, leading eventually to a constant efflux/influx rate regardless of the exact auxin concentration. Note also that production and decay of auxin have been set to 1 by locking in our choice for units of time and auxin concentration.

From this expression, we can also observe the contribution to outgoing active auxin flows by the term with $p_{i\,i+1}$. It makes sense to argue that for each wall, the direction of this flow is perpendicular to the wall, $\mathbf{n}_{i\,i+1}$, pointing outwards. Were one to track the auxin flow as a vector, it would result in,

$$\overrightarrow{\text{active auxin outflow}}_\alpha = \frac{P_\alpha}{A_\alpha}\frac{a_\alpha}{K_{\text{efflux}} + a_\alpha}\sum_{i\in\mathcal{V}_\alpha} l_{i\,i+1}p_{i\,i+1}\mathbf{n}_{i\,i+1} \propto \sum_{i\in\mathcal{V}_\alpha} l_{i\,i+1}p_{i\,i+1}\mathbf{n}_{i\,i+1} = \mathbf{F}_\alpha. \tag{13}$$

Active auxin outflow is therefore proportional to the polarity vector $\mathbf{F}_\alpha$.

Notoriously, this description lacks diffusion of auxin across cell wall compartments and, therefore, apoplastic pathways of auxin transport, which is a significant contribution [16]. Notwithstanding, and according to [25], this approximation seems valid in a plethora of plant tissues, and it can reproduce the patterns found therein. Here, we use a similar approach to avoid compounding further complexity onto the model. For the same reason, another approximation implicit in the model used is the lack of internal auxin gradients that exist in elongated cells [16, 18].

##### 1.4.2 Mechanical regulation of efflux carriers

Although $q_{i\,i+1}$ will be set according to experimental distributions, $p_{i\,i+1}$ will be computed from the steady state of a parsimonious carrier binding model, also in accordance with [7]. Effectively, we use

$$p_{i\,i+1} = \frac{f_{i\,i+1}}{1 + \sum_{k\in\mathcal{V}_\alpha}\frac{l_{k\,k+1}}{L_\alpha}f_{k\,k+1}}, i \in \mathcal{V}_\alpha, \tag{14}$$

where $f_{i\,j}$ denotes the ratio between carrier binding and unbinding rates associated with the same compartment as $p_{i\,j}$. As per the hypothesis explored throughout this work, we implement mechanical regulation by writing this ratio as

$$f_{i\,i+1} = \begin{cases}\left(\eta\sigma_{i\,i+1}\right)^n, & \sigma_{i\,i+1} > 0\\ 0, & \sigma_{i\,i+1} \leq 0\end{cases}, i \in \mathcal{V}_\alpha, \tag{15}$$

where $\eta$ and $n$ are parameters that quantify stress perception and non-linear response, respectively, and $\sigma_{i\,j}$ is the value of longitudinal stress acting on this specific compartment. Given an optimized geometry, using current wall lengths, $l_{i\,j}$, rest wall lengths, $l_{0\,i\,j}$, and cell wall stiffness $E_\alpha$, the stress acting on a wall of cell $\alpha$ can simply written down as $\sigma_{i\,j} = E_\alpha\left(l_{i\,i+1} - l_{0\,i\,i+1}\right)/l_{0\,i\,i+1}$, $i \in \mathcal{V}_\alpha$.

This steady-state solution describes how exporters would be distributed for constant auxin distribution for a long time. Since our particular system has a stable auxin maximum with only small perturbing effects, this approximation is valid.

##### 1.4.3 Auxin-mediated cell wall softening

As cells grow, auxin plays a role in disrupting the cell wall to allow it to flow and elongate [23, 4]. As such, and taking inspiration from the same work [7], we model auxin-mediated cell wall softening as auxin-dependent variable stiffness,

$$E_\alpha = E\left(a_\alpha\right) = E_0 \left(1 + r \frac{1 - a_\alpha^m}{1 + a_\alpha^m}\right), \tag{16}$$

where $E_0$ represents the value of stiffness for basal levels of auxin, $a_\alpha = 1$, the softening parameter, $r$, is contained in $[0, 1[$, such that $E_\alpha$ is confined to the interval $]E_0\left(1 - r\right), E_0\left(1 + r\right)[$, and $m$ characterizes the non-linear response of auxin-mediated cell wall softening.

### 2 Parametrization

The length scale ($L = 1$) is set to the width of the pericycle layer prior to pressurization. We maintain the same scale for height, implying a rectangular prism geometry with isotropic base shape. With a width of $w = 0.08$, the cell wall thickness is estimated to be of the order of $\sim 100\,\text{nm}$.

Mechanical regulation parameters $(n, r, m, \eta)$ are sourced from [9, 7]. Turgor pressure ($T = 0.65\,\text{MPa}$) is based on measurements from [2], while the base elastic modulus ($E_0 = 300\,\text{MPa}$) is taken from [28]. The increased elastic modulus in the vasculature is assumed due to lignin content in the xylem [10].

Auxin transport parameters $K_{\text{efflux}}$ and $K_{\text{influx}}$ are assumed, such that they saturate activity for the same value of auxin for both carriers. The value chosen reflects that of [9].

Time and auxin concentration units are normalized to 1. Values for metabolism, $P$, $Q$, and $D$ are estimated from [17] and [24] respectively, but adjustments are made to achieve a stable auxin maximum in founder cells. Specifically, $P$ is decreased and $Q$ and $D$ are increased, resulting in an underestimation of efflux carrier potency.

The bending modulus ($k_b = 3600\,\text{MPa}$) is chosen to match tissue geometry (Fig. S5), validated by blocking radial expansion of founder cells as observed when the endodermis restricts growth [27].

Extensibility is relatively large when compared to the auxin turnover rate, but further reduction is hindered by simulation time constraints.

| parameter | value | parameter | value |
|---|---|---|---|
| height | $h = 1$ | width | $w = 0.08$ |
| default bending modulus | $k_b = 3600\,\mathrm{MPa}$ | default turgor | $T = 0.65\,\mathrm{MPa}$ |
| auxin production rate | 1 | auxin decay rate | 1 |
| diffusion strength | $D = 7200$ | active efflux strength | $P = 1800$ |
| active inflow strength | $Q = 36000$ | efflux saturation constant | $K_{\mathrm{efflux}} = 1$ |
| influx saturation constant | $K_{\mathrm{influx}} = 1$ | stress perception constant | $\eta = 10\,\mathrm{MPa}^{-1}$ |
| stress perception exponent | $n = 3$ | base elastic modulus | $E_0 = 300\,\mathrm{MPa}$ |
| softening parameter | $r = 2/3$ | softening exponent | $m = 2$ |
| founder cells extensibility | $\varphi = 0.05\,\mathrm{MPa}^{-1}$ | default extensibility | $\varphi = 0\,\mathrm{MPa}^{-1}$ |
| vasculature base elastic modulus | $E_0 = 3000\,\mathrm{MPa}$ | | |

Table S1: **Values of parameters used.** Only the parameters relating to stress have units. All other unit measures have been normalized. For instance, length is in units of the pericycle width at rest before growth. Units of time and auxin concentration are such that production and decay are 1, meaning units of concentration are relative to a basal auxin concentration, $a_0$, which takes time $t = 1$ to reach $(1 - 1/e)\,a_0$ from 0 in an isolated cell.

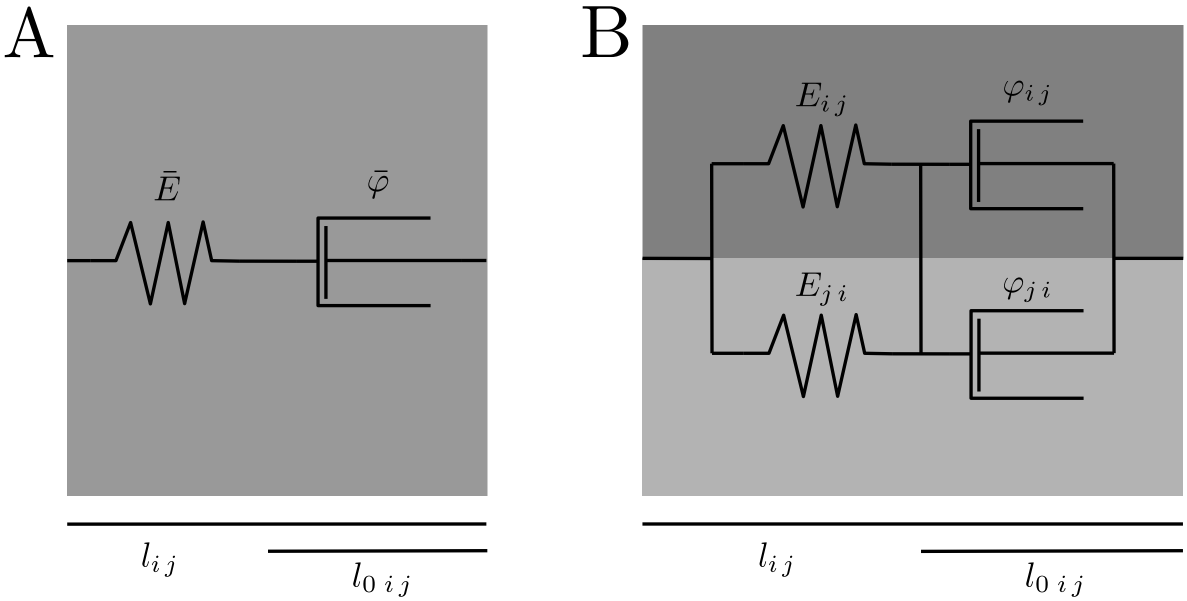


Figure S1: **Rheological models used in this work.** (A) Rheological model of a regular Maxwell material. It is the same constitutive model used by [20, 5]. (B) Our modified version to account for two wall compartments with different mechanical parameters while keeping rest length, $l_0$, the same between compartments. We assume each viscous element is under the stress corresponding to the same wall compartment. The model in (B), under this assumption, admits an equivalent model, (A), when associating elastic moduli, $E_{i\,j}$ and extensibilities, $\varphi_{i\,j}$, according to Eq. 11.

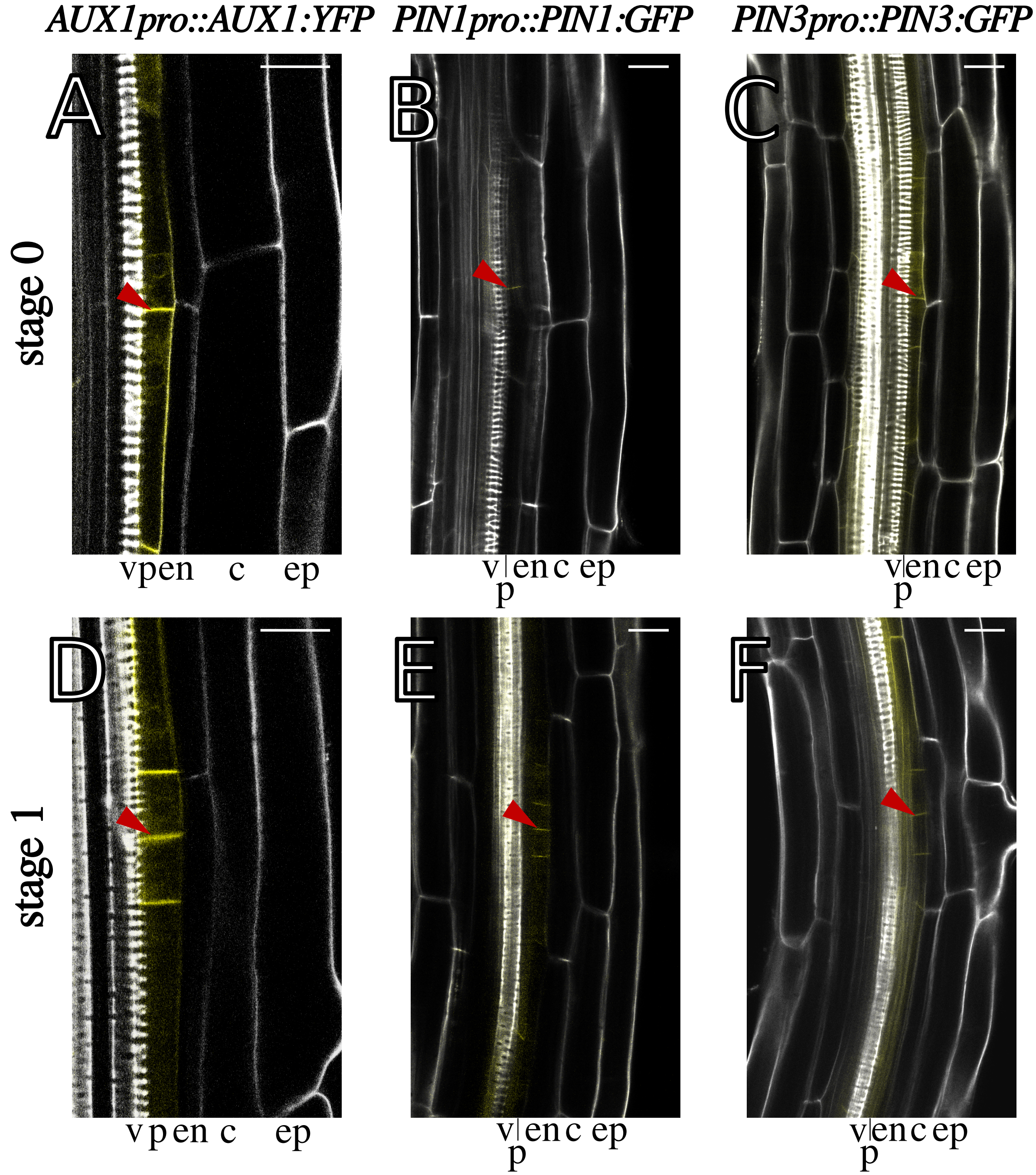


Figure S2: **Comparison of carrier distributions (A-C) before (stage 0) and (D-F) after LR radial expansion (stage 1).** (A and D) Influx carriers of the AUX1 family keep their activity exclusive to the activated XPP cells, particularly their daughter cells. (B, C, E, F) Efflux carriers of both families have similar behaviour in that they are present in and around the activated XPP cells. In particular, efflux carriers of the PIN1 family seem to prefer binding to the shared interface between activated XPP cells (red arrowhead). Moreover, efflux carriers do not vanish after stage 0, which, according to our model, would indicate that stress would not have decayed completely. The different layers are labelled: vasculature (v), pericycle (p), endodermis (en), cortex (c). Scale bar: 20 $\mu$m.

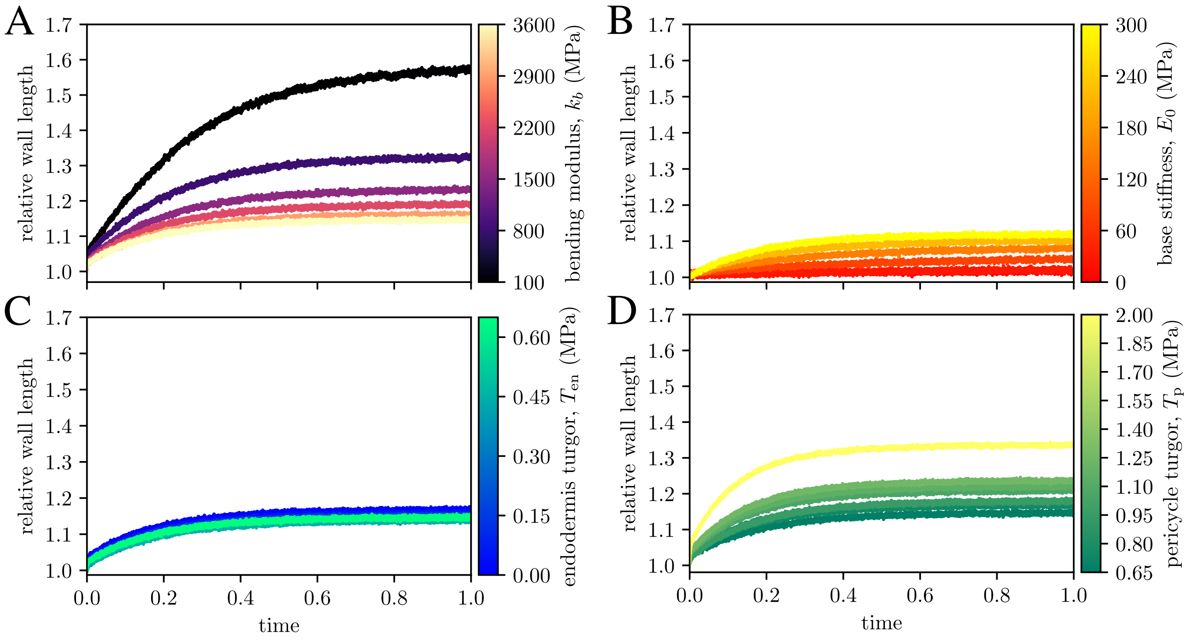


Figure S3: Related to Figure 4. **Relative wall length of the anticlinal wall shared between activated pericycle cells as a function of time and of mechanical parameters**. (A) local bending stiffness, $k_b$, (B) local stiffness, $E_0$, (C) overlaying endodermis turgor, $T_{\mathrm{en}}$, and (D) activated pericycle turgor, $T_{\mathrm{p}}$.

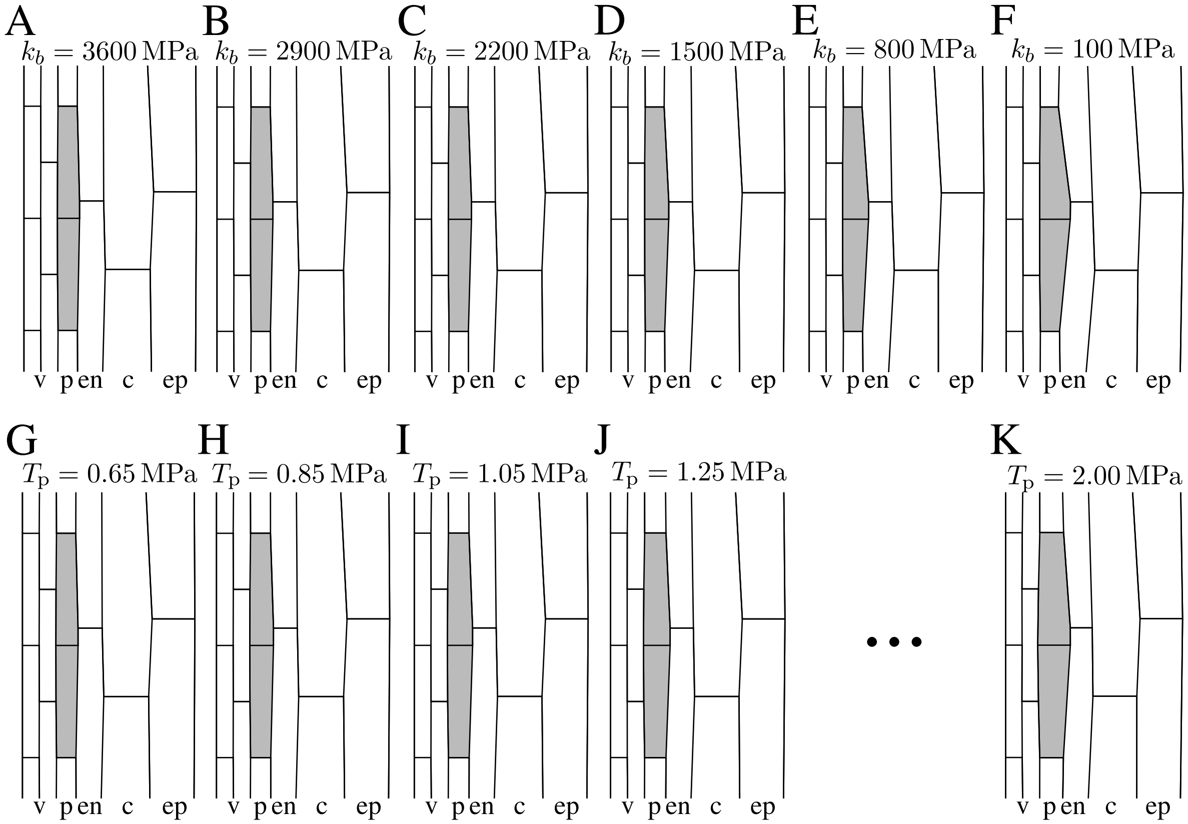


Figure S4: Related to Figure 4. **Activated pericycle cells shape as a function of mechanical parameters**. (A-F) Decreasing bending modulus, $k_b$. (G-K) Increasing primed pericycle turgor, $T_p$. The values for the respective parameters presented are (A) $k_b = 3600\,\mathrm{MPa}$, (B) $k_b = 2900\,\mathrm{MPa}$, (C) $k_b = 2200\,\mathrm{MPa}$, (D) $k_b = 1500\,\mathrm{MPa}$, (E) $k_b = 800\,\mathrm{MPa}$, (F) $k_b = 100\,\mathrm{MPa}$, (G) $T_p = 0.65\,\mathrm{MPa}$, (H) $T_p = 0.85\,\mathrm{MPa}$, (I) $T_p = 1.05\,\mathrm{MPa}$, (J) $T_p = 1.25\,\mathrm{MPa}$, (K) $T_p = 2.00\,\mathrm{MPa}$. Note that (A) and (G) are both the same control experiment. The labelled layers are vasculature (v), pericycle (p), endodermis (en), cortex (c), and epidermis (ep).

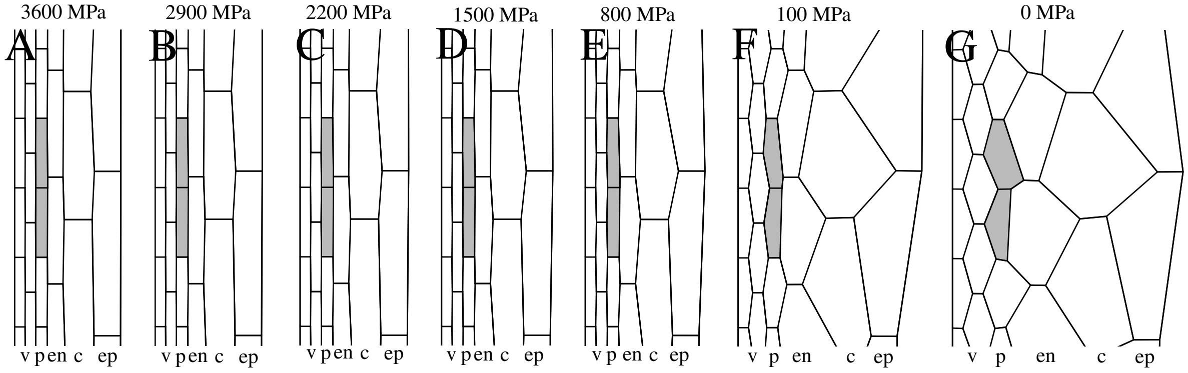


Figure S5: Related to Figure 4. **Geometry of cell template for different bending stiffness values,** $k_b$. The labelled layers are vasculature (v), pericycle (p), endodermis (en), cortex (c), and epidermis (ep).

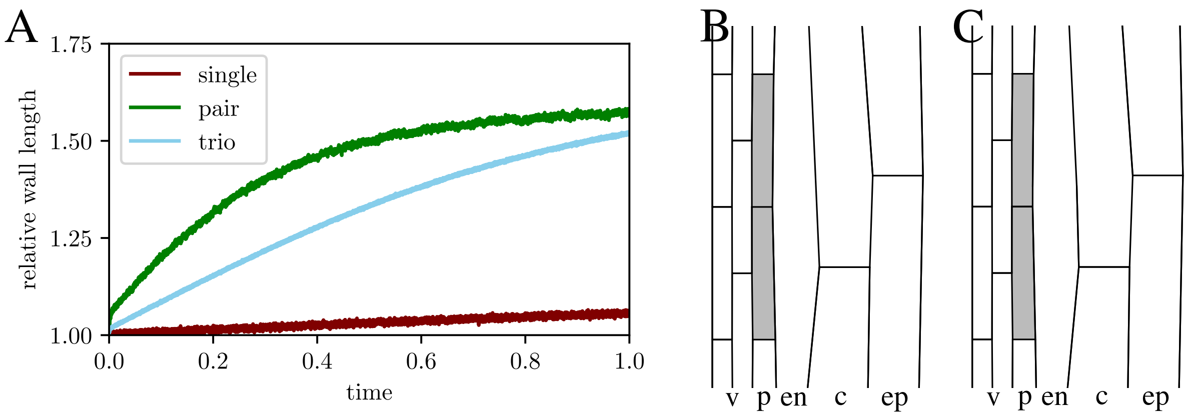


Figure S6: Related to Figure 4. **Relative wall length over time for a different number of overlying endodermal cells**. These correspond to the final states presented in Fig. 4 H-J in the main text. Final states, at $t = 1$ for a single overlaying endodermal cell where (B) junctions where the wall was previously were removed, and (C) where the junctions were left after the removal of the wall and are free to move during optimization. The labelled layers are vasculature (v), pericycle (p), endodermis (en), cortex (c), and epidermis (ep).

### References

[1] S. Alt, P. Ganguly, and G. Salbreux. Vertex models: from cell mechanics to tissue morphogenesis. *Philosophical Transactions of the Royal Society B: Biological Sciences*, 372(1720):20150520, may 2017.

[2] L. Beauzamy, M. Louveaux, O. Hamant, and A. Boudaoud. Mechanically, the shoot apical meristem of arabidopsis behaves like a shell inflated by a pressure of about 1 MPa. *Frontiers in Plant Science*, 6(NOVEMBER):1–10, 2015.

[3] E. Benková, M. Michniewicz, M. Sauer, T. Teichmann, D. Seifertová, G. Jürgens, and J. Friml. Local, Efflux-Dependent Auxin Gradients as a Common Module for Plant Organ Formation. *Cell*, 115(5):591–602, nov 2003.

[4] S. A. Braybrook and A. Peaucelle. Mechano-Chemical Aspects of Organ Formation in Arabidopsis thaliana: The Relationship between Auxin and Pectin. *PLoS ONE*, 8(3):e57813, mar 2013.

[5] F. Corson, O. Hamant, S. Bohn, J. Traas, A. Boudaoud, and Y. Couder. Turning a plant tissue into a living cell froth through isotropic growth. *Proceedings of the National Academy of Sciences USA*, 106(21):8453–8458, may 2009.

[6] J. Cufí, A. Reventós, and C. J. Rodríguez. Curvature for polygons. *The American Mathematical Monthly*, 122(4):332, 2015.

[7] D. Ramos, João R., Maizel, Alexis, and Alim, Karen. Tissue-wide integration of mechanical cues promotes effective auxin patterning. *Eur. Phys. J. Plus*, 136(2):250, 2021.

[8] H. Deuling and W. Helfrich. Red blood cell shapes as explained on the basis of curvature elasticity. *Biophysical Journal*, 16(8):861–868, 1976.

[9] M. G. Heisler, O. Hamant, P. Krupinski, M. Uyttewaal, C. Ohno, H. Jönsson, J. Traas, and E. M. Meyerowitz. Alignment between PIN1 Polarity and Microtubule Orientation in the Shoot Apical Meristem Reveals a Tight Coupling between Morphogenesis and Auxin Transport. *PLoS Biology*, 8(10):e1000516, oct 2010.

[10] D. Hepworth and J. Vincent. The mechanical properties of xylem tissue from tobacco plants (nicotiana tabacum‘samsun’). *Annals of Botany*, 81(6):751–759, 1998.

[11] H. Jonsson, M. G. Heisler, B. E. Shapiro, E. M. Meyerowitz, and E. Mjolsness. An auxin-driven polarized transport model for phyllotaxis. *Proceedings of the National Academy of Sciences USA*, 103(5):1633–1638, jan 2006.

[12] J.-D. Julien, A. Pumir, and A. Boudaoud. Strain- or stress-sensing in mechanochemical patterning by the phytohormone auxin. *bioRxiv*, 2019.

[13] J. Khadka, J.-D. Julien, and K. Alim. Feedback from tissue mechanics self-organizes efficient outgrowth of plant organ. *Biophysical Journal*, 117(10):1995–2004, nov 2019.

[14] J. Kierfeld, K. Baczynski, P. Gutjahr, and R. Lipowsky. Semiflexible polymers and filaments: From variational problems to fluctuations. *AIP Conference Proceedings*, 1002:151, 04 2008.

[15] J. Kierfeld, O. Niamploy, V. Sa-yakanit, and R. Lipowsky. Stretching of semiflexible polymers with elastic bonds. *The European Physical Journal E*, 14(1):17–34, May 2004.

[16] E. M. Kramer. Computer models of auxin transport: a review and commentary. *Journal of Experimental Botany*, 59(1):45–53, 04 2007.

[17] E. M. Kramer and E. M. Ackelsberg. Auxin metabolism rates and implications for plant development. *Frontiers in Plant Science*, 6(March):1–8, mar 2015.

[18] O. Leyser. Auxin Signaling. *Plant Physiology*, 176(1):465–479, jan 2018.

[19] L. Livadaru, R. R. Netz, and H. J. Kreuzer. Stretching response of discrete semiflexible polymers. *Macromolecules*, 36(10):3732–3744, Apr. 2003.

[20] J. A. Lockhart. An analysis of irreversible plant cell elongation. *Journal of Theoretical Biology*, 8(2):264–275, mar 1965.

[21] Y. Long, I. Cheddadi, G. Mosca, V. Mirabet, M. Dumond, A. Kiss, J. Traas, C. Godin, and A. Boudaoud. Cellular heterogeneity in pressure and growth emerges from tissue topology and geometry. *Current Biology*, 30(8):1504 – 1516.e8, 2020.

[22] J. K. E. Ortega. Augmented Growth Equation for Cell Wall Expansion. *Plant Physiology*, 79(1):318–320, sep 1985.

[23] A. Peaucelle, S. A. Braybrook, L. Le Guillou, E. Bron, C. Kuhlemeier, and H. Höfte. Pectin-Induced Changes in Cell Wall Mechanics Underlie Organ Initiation in Arabidopsis. *Current Biology*, 21(20):1720–1726, oct 2011.

[24] H. L. Rutschow, T. I. Baskin, and E. M. Kramer. The carrier AUXIN RESISTANT (AUX1) dominates auxin flux into Arabidopsis protoplasts. *New Phytologist*, 204(3):536–544, nov 2014.

[25] P. Sahlin, B. Söderberg, and H. Jönsson. Regulated transport as a mechanism for pattern generation: Capabilities for phyllotaxis and beyond. *Journal of Theoretical Biology*, 258(1):60–70, may 2009.

[26] R. S. Smith, S. Guyomarc'h, T. Mandel, D. Reinhardt, C. Kuhlemeier, and P. Prusinkiewicz. A plausible model of phyllotaxis. *Proceedings of the National Academy of Sciences USA*, 103(5):1301–1306, jan 2006.

[27] J. E. M. Vermeer, D. von Wangenheim, M. Barberon, Y. Lee, E. H. K. Stelzer, A. Maizel, and N. Geldner. A Spatial Accommodation by Neighboring Cells Is Required for Organ Initiation in Arabidopsis. *Science*, 343(6167):178–183, jan 2014.

[28] A. Weber, S. Braybrook, M. Huflejt, G. Mosca, A.-L. Routier-Kierzkowska, and R. S. Smith. Measuring the mechanical properties of plant cells by combining micro-indentation with osmotic treatments. *Journal of Experimental Botany*, 66(11):3229–3241, jun 2015.
